# Retrograde mitochondrial transport is essential for mitochondrial homeostasis in neurons

**DOI:** 10.1101/683243

**Authors:** Amrita Mandal, Katherine Pinter, Natalie Mosqueda, Alisha Beirl, Richa Madan Lomash, Sehoon Won, Katie S Kindt, Catherine M Drerup

## Abstract

Mitochondrial transport in neurons is essential for forming and maintaining axonal projections. While much is known about anterograde mitochondrial movement, the function of retrograde mitochondrial motility in neurons was unknown. We investigated the dynamics and utility of retrograde mitochondrial transport. Using long-term tracking of mitochondria in vivo, we found mitochondria in axon terminals turnover within hours via retrograde transport. Mitochondria do not return to the cell body solely for degradation; rather, mitochondria use bidirectional transport to redistribute themselves throughout the neuron. Disruption of retrograde mitochondrial transport severely depletes the cell body of mitochondria and impacts mitochondrial health throughout the cell. Altered mitochondrial health correlates with decreased synaptic activity. Using proteomics, we provide evidence that retrograde mitochondrial movement functions to maintain the organelle’s proteome. Together, our work demonstrates that mitochondrial retrograde transport is essential for the maintenance of a homeostatic population of mitochondria in neurons and consequently effective synaptic activity through promoting mitochondrial protein turnover.

## Introduction

Mitochondria are essential for ATP production and play vital roles in other cellular processes including acting as signaling centers, regulating local calcium levels, and serving as the primary storage and utilization site for iron in the cell ^1–7^. Not surprisingly given these diverse and critical roles, impaired mitochondrial function can lead to a number of disease states. In neurons, mitochondria have been shown to localize to synapses to maintain local ATP levels essential for synaptic activity ^8,9^. Additionally, this organelle participates in the active buffering of calcium in the axon terminal which modulates synaptic release ^10–12^. More recent work has shown that mitochondria also fuel protein production in distal neuronal compartments, implicating this organelle in the regulation of local translation ^13,14^. Therefore, maintaining a healthy pool of mitochondria, particularly in distal compartments of the neuron, is critical for the health and maintenance of functional neural circuits. While much is known about how damaged mitochondria are cleared from this region ^15–17^, less is known about how populations of healthy mitochondria are maintained in neurons over their lifetime.

Mitochondrial maintenance is complicated by the fact that this organelle requires more than a thousand proteins for optimal health and function. While mitochondria maintain their own genome ^18–20^, which includes genes encoding 13 proteins in humans, the bulk of the proteins important for the function and maintenance of this organelle are synthesized from genes encoded in the nucleus ^21,22^. These proteins have diverse half-lives, ranging from hours to weeks ^23,24^. Once generated, mitochondrial proteins translocate to the correct compartment within the organelle through well-described mitochondrial protein import pathways ^25^. However, while we know how proteins are incorporated into the organelle, how they are brought to the organelle prior to import, particularly in distal compartments, is largely unknown. While transport of proteins and/or mRNAs could be sufficient to replenish a number of mitochondrial proteins, it would require active transport of individual components to organelles which can be a meter from the cell body in humans. Given the number of mitochondrial proteins, the rapid turnover rates of a subset of them, and the distance from the cell body to distal neuronal compartments, active transport of each individual protein would be energy intensive ^24^. Alternatively, mitochondria themselves could be transported from the distal processes to the cell body for protein turnover. Currently, there is little evidence to support either of these models.

Mitochondria move throughout the neuron using the Kinesin-1 and Cytoplasmic dynein motors for anterograde and retrograde transport respectively ^26–29^. While mitochondrial transport is frequent during axon extension, work in vitro and in vivo has shown a substantial decrease in mitochondrial movements in mature axons ^11,30^. Perturbation of mitochondrial function in mature axons leads to rapid loss of mitochondrial health which is correlated with varying degrees of enhanced retrograde mitochondrial movement ^16,31,32^. This is controversial, however, as others have shown that mild mitochondrial stress, which still impacts organelle health, halts all movement ^15,33^. Additionally, mitochondrial membrane potential (correlate of health) does not correlate with direction of transport ^34^. Much of this work on mitochondrial transport with and without mitochondrial damage has been done on the order of minutes. Significantly less is known about mitochondrial movement that occurs over longer time scales (hours to days) and the real time dynamics and life time of this organelle in non-pathological states. This gap in our knowledge is due to the lack of an in vivo system that can be monitored at a cellular level for longer periods of time. Zebrafish offer the advantages of larval transparency and easily accessible functional neural circuits in which we can image the movement, health, and function of mitochondria in vivo in normative conditions over several days.

Using live imaging of mitochondrial dynamics in vivo, we show that mitochondria in mature neurons move in the retrograde direction quite frequently over the course of hours. This retrograde movement is essential to maintain a substantial portion of the mitochondrial proteome in the distal axon. When retrograde mitochondrial movement is disrupted, ∼30% of mitochondrial proteins show a greater than 50% reduction. Supporting the necessity of retrograde mitochondrial movement in axons, inhibition of this process severely disrupts the neuronal mitochondrial population: Mitochondria are lost from the cell body and accumulate in axon terminals. When stationary mitochondria accumulate in distal axonal compartments, they show evidence of failing mitochondrial health including: elevated levels of chronic reactive oxygen species, reduced matrix potential, and decreased calcium load. Finally, synapses in neurons with inhibited retrograde mitochondrial transport show significantly diminished activity. Together, our work argues that mitochondria move in the retrograde direction in neurons to reposition themselves near the primary site of protein production in the cell body. This movement is essential to replenish critical proteins essential for organelle and neural circuit health and function.

## Results

### Mitochondrial populations turnover within 24 hrs in neuronal compartments

Imaging of neuronal mitochondrial dynamics has typically been done on the order of minutes in cultured neurons and more recently in vivo in mouse brain and sciatic nerve ^30,35^. This work demonstrated that mitochondrial movement in mature axons is infrequent over several minutes; however, the frequency of mitochondrial movement in axons over hours has not been analyzed. To directly test whether mitochondria are static in mature axons over hours and days, we engineered a transgenic zebrafish (*Tg(5kbneurod:mito-mEos)*^*y568*^) to label mitochondria in neurons with the photoconvertible protein, mEos ^36^. This transgene labels many neurons, including the afferent axons and soma of posterior lateral line (pLL) neurons (Fig. 1a-c). We used the pLL neurons to analyze mitochondrial localization and turnover because their superficial localization enables straightforward in vivo imaging ^37^. The pLL is a mechanosensory system in the zebrafish. Afferent axons extend from the sensory neuron cell bodies in the pLL ganglion and innervate sensory hair cells (HCs) in organs called neuromasts (NMs) situated along the trunk (reviewed in ^38–40^). This sensory system is fully formed and functional by 4 days post-fertilization (dpf), which is the earliest time-point analyzed in the work below.

**Figure 1:**
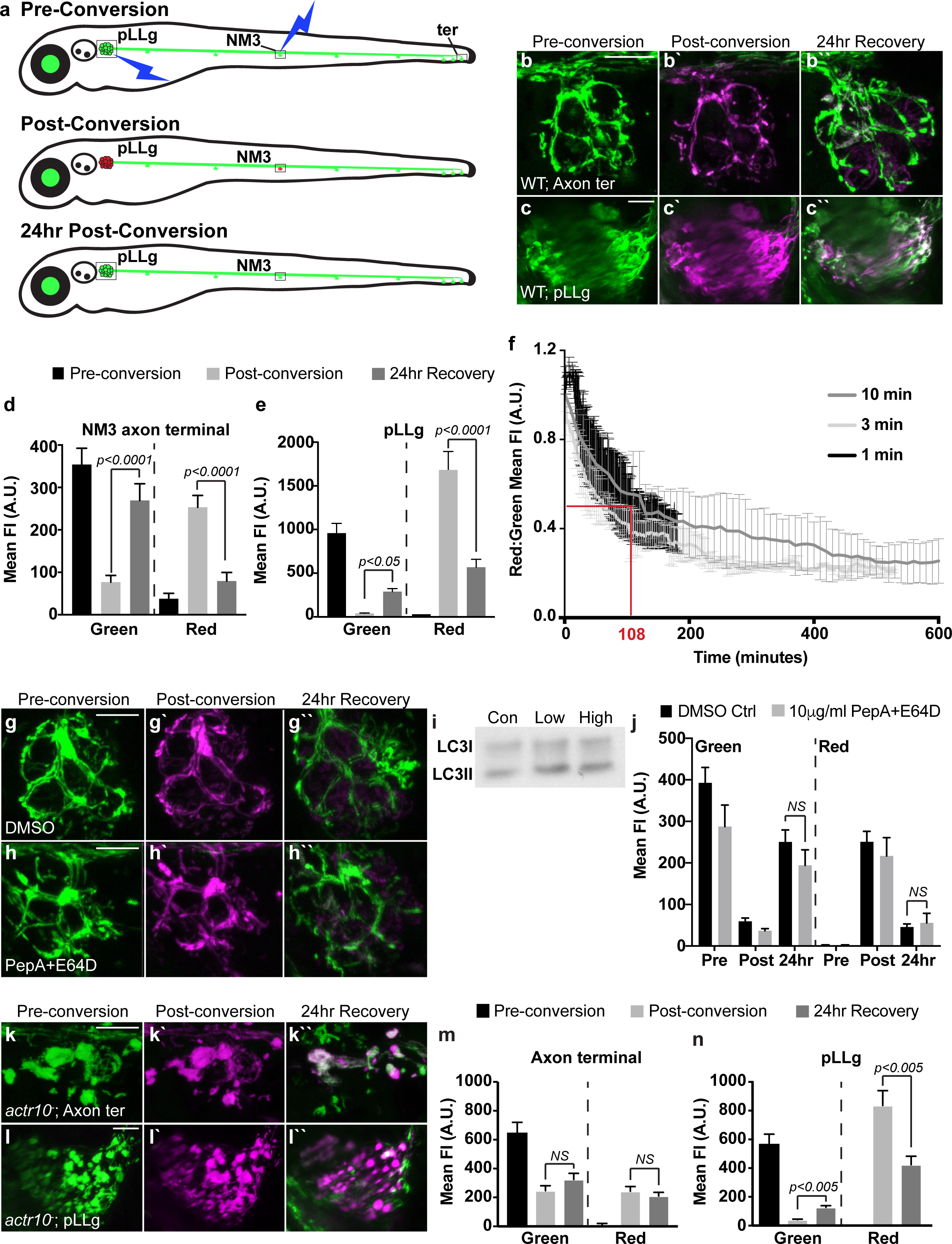
Active mitochondrial transport is required for mitochondrial turnover in neuronal compartments. (**a**) Schematic of mitochondrial photoconversion assay using the *Tg(5kbneurod:mito-mEos)* transgenic zebrafish. Sensory neurons of the posterior lateral line ganglion (pLLg) and axon terminals of neuromast 3 (NM3) or the terminal cluster (ter) are depicted. (**b**,**c**) Photoconversion of axon terminals (**b**) and the cell bodies in the ganglion (**c**) results in a permanent switch from green to red (shown in magenta) of the mitochondrially localized mEos (photoconversion at 4 days post-fertilization (dpf)). These converted mitochondria are gone 24 hrs after photoconversion. (**d**,**e**) Quantification of the gain of new (green) and loss of old (magenta) mitochondria from the axon terminal (**d**; n=7) or pLLg (**e**; n=18) 24 hrs after photoconversion (ANOVA; Tukey HSD posthoc contrasts). (**f**) Time-lapse imaging of mitochondrial turnover in axon terminals reveals that 50% of mitochondria have left the axon terminal by 108 minutes post-conversion (1 min: n=4; 3 min:n=2; 10 min:n=2). (**g**-**j**) Mitochondrial turnover in axon terminals is not affected by lysosomal inhibition. (**i**) Western blot of whole embryo extracts after treatment with Pepstatin A and E64D (2μg/mL-low; 10μg/mL-high for ∼18hrs) reveals increased LC3II. (**j**) Quantification of mitochondrial turnover with lysosomal inhibition (ANOVA; n=11; NS – not significant). (**k**-**n**) Retrograde mitochondrial transport is necessary for mitochondrial turnover. (**k**,**l**) Photoconversion of mitochondrially-localized mEos in axon terminals and the pLLg of *actr10*^*-*^ mutants. (**m**,**n**) Quantification of new (green) and old (magenta) mitochondria shows persistence of old converted mitochondria in axon terminals, when retrograde transport is disrupted. pLLg mitochondrial turnover is intact. (ANOVA; Tukey HSD posthoc contrasts; n=7). Scale bars – 10μm.

We first used this system to examine mitochondrial turnover in the neuronal cell bodies and axon terminals. We either photoconverted the mitochondrially localized mEos in a mid-trunk axon terminal (NM3) or in the cell bodies of the pLL ganglion (pLLg; Fig. 1a-c). We then measured green (native) and red (photoconverted) mEos fluorescence intensities in the converted compartment immediately after conversion and 24 hrs later. Quantification of fluorescence intensities revealed addition of new (green) mitochondria and loss of the photoconverted (red) mitochondria from the cell bodies by 24 hrs post-conversion (Fig. 1c,e), likely due to anterograde transport of organelles from this region into the axon. Somewhat surprisingly, we also saw significant loss of axon terminal mitochondria in this time-period, indicating almost complete organelle turnover within a day (Fig. 1b,d). Similar trends were obtained in all experiments performed for axon terminals: 1) in mature mid-trunk axons (4-5 dpf; Fig. 1); 2) in 4-5 dpf axons that innervate the distal trunk (ter); 3) in mid-trunk and distal trunk axons at later stages (6-7 dpf) when myelination of the nerve is complete (Supplementary Fig. 1a-c); and 4) in primary motor neuron axons (Supplementary Fig. 1f-h).

To determine the precise temporal dynamics of this turnover, we performed rapid time-lapse imaging of axon terminals immediately after photoconversion. These time-lapse imaging experiments revealed that 50% of the converted mitochondria are depleted from the axon terminal 108 minutes after conversion (Fig. 1f). For both the time-lapse and 24 hr interval analyses, larvae were immobilized in low-melt agarose for the duration of the experiment. To control for any effects of mounting on mitochondrial turnover, we performed the 24 hr interval experiments at 4-5 dpf and 6-7 dpf but removed the larvae from the agarose enclosures between imaging sessions. Similar addition of new and loss of old mitochondria was apparent from axon terminals with these conditions (Supplementary Fig. 1d,e). Together, our data support reliable mitochondrial turnover in mature sensory and motor neuron axons on the scale of hours.

### Mitochondria redistribute from axon terminals throughout the neuron via retrograde transport

Mitochondrial turnover in neuronal compartments could be due to local degradation or retrograde transport. To determine if lysosomal degradation contributed to loss of mitochondria from axon terminals, we photoconverted mitochondria marked by mEos and then treated with lysosomal inhibitors Pepstatin A and E64D (Fig. 1g,h)^41,42^. Western analysis of whole larval extracts showed elevation of LC3II, indicating inhibition of overall lysosomal degradation; however, no change in the turnover of mitochondria in axon terminals following lysosomal inhibition was observed (Fig. 1i,j). Next, we assessed whether retrograde mitochondrial transport could account for the turnover of mitochondria from axon terminals. For this, we utilized a zebrafish mutant strain, *actr10*^*nl15*^ (hereafter referred to as *actr10*^*-*^), shown to specifically disrupt the retrograde transport of mitochondria ^43^. Analysis of mitochondrial persistence in this line revealed no change in red (old) mitochondria 24 hr after photoconversion in axon terminals (Fig. 1k,m). Thus, retrograde transport is the primary mediator of mitochondrial turnover in axon terminals. Photoconversion of the pLL ganglion demonstrated similar mitochondrial turnover between *actr10*^*-*^ mutants and wildtype animals, supporting intact anterograde transport in this line as has been previously described (Fig. 1l,n) ^43^.

To determine where mitochondria go after they are retrogradely transported from the axon terminal, we photoconverted mitochondria in two axon terminals of the mid-trunk sensory organ (NM3) and 24 hrs later imaged four distinct regions within the converted neurons: 1) the neuronal cell bodies in the pLLg; 2) axons in the proximal pLL nerve; 3) a region of the pLL nerve adjacent to the photoconverted terminals; and 4) a region of the pLL nerve distal to the axon terminals that were converted (negative control; Fig. 2i). Immediately following photoconversion, we observed a few converted (red) mitochondria in the nerve adjacent to the axon terminal. These organelles were likely actively moving from the terminal during conversion. The cell bodies and proximal axon were devoid of converted organelles at this time-point (Fig. 2a-d). 24 hrs after conversion, mitochondria were visible in the 2 converted axons in all regions of the pLL nerve proximal to the converted terminal as well as in 2 cell bodies of the pLLg (Fig. 2e-h, j). This data indicates that mitochondria are retrogradely transported from axon terminals and redistributed throughout the neuron over the course of a day.

**Figure 2:**
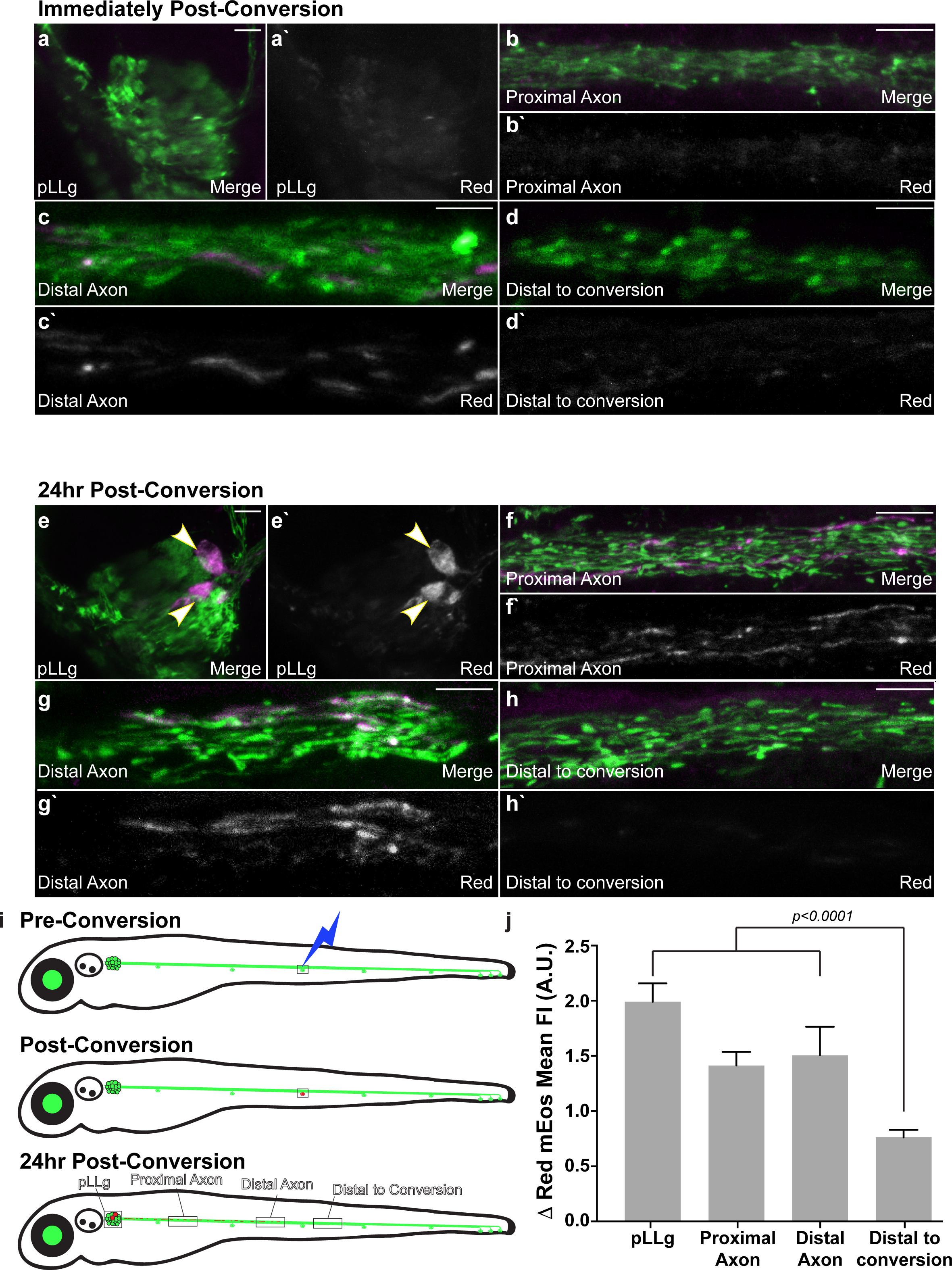
Mitochondrial retrograde transport results in redistribution of axon terminal mitochondrial throughout the neuron. (**a**-**d**) Immediately after photoconversion, converted mitochondria are only detected immediately proximal to the converted axon terminal (distal axon; shown in magenta above and white below). (**e**-**h**) 24 hrs after photoconversion, mitochondria that originated from the converted axon terminal are now redistributed to the neuronal cell bodies (**e**), proximal axons (**f**), and distal axons (**g**). (**h**) Axons distal to the converted terminals do not contain converted mitochondria. (**i**) Schematic of the photoconversion and areas imaged. (**j**) Quantification of the change in magenta (converted) fluorescence intensity 24 hrs after photoconversion (n=12; ANOVA; Tukey HSD posthoc contrasts). Scale bars – 10μm.

### Inhibition of retrograde mitochondrial transport leads to an imbalance in neuronal mitochondrial load

Our analysis of mitochondrial localization after photoconversion demonstrated that mitochondria redistribute from axon terminals throughout the neuron using retrograde transport. We next assayed the impact of disrupted mitochondrial retrograde transport on mitochondrial localization and health in neurons. To study localization, we utilized a sparse labeling approach in *actr10*^*-*^ mutant zebrafish ^43^. Injection of the *5kbneurod:mito-TagRFP* DNA plasmid into zygotes leads to mosaic expression of mitochondrially localized TagRFP in a subset of pLL neurons ^36,37^. We used this method to image mitochondrial load (mitochondrial area/neuronal area) in the soma, proximal axon, distal axon, and axon terminal of neurons (Fig. 3a). While mitochondria fill the cell body of a wildtype neuron, inhibition of retrograde mitochondrial movement leads to a significant reduction of mitochondrial load in the soma and an accumulation of this organelle in axon terminals both at 4 and 6 dpf (Fig. 3b,c,e-j,l,m). To confirm this altered distribution was due to disrupted retrograde mitochondrial movement, we also analyzed mitochondrial localization in a mutant with loss of function mutations in both zebrafish paralogs of p150 (*dctn1a/dctn1b*), a core subunit of the dynein accessory complex dynactin ^44^. Loss of p150 leads to cessation of the majority of retrograde cargo motility, including that of mitochondria. *p150a*^*-*^*/b*^*-*^ double mutants had a reduction of mitochondrial load in the cell body and accumulation in axon terminals similar to *actr10*^*-*^ mutants (Fig. 3d,k,l). Finally, we wanted to determine if inhibiting mitochondrial retrograde transport in mammalian neurons would also alter mitochondrial load in the cell body. For this analysis, we knocked down Actr10 in rat hippocampal neurons using shRNA. Testing of three targets revealed that shRNA #2 (see Resources table) was the most effective (Fig. 3p,q). Knock down of Actr10 in rat hippocampal neurons significantly reduced cell body mitochondrial load, similar to what was observed in zebrafish neurons (Fig. 3n,o,r). Together, this data supports a role for active retrograde mitochondrial transport in the maintenance of a homeostatic distribution of mitochondria in neurons.

**Figure 3:**
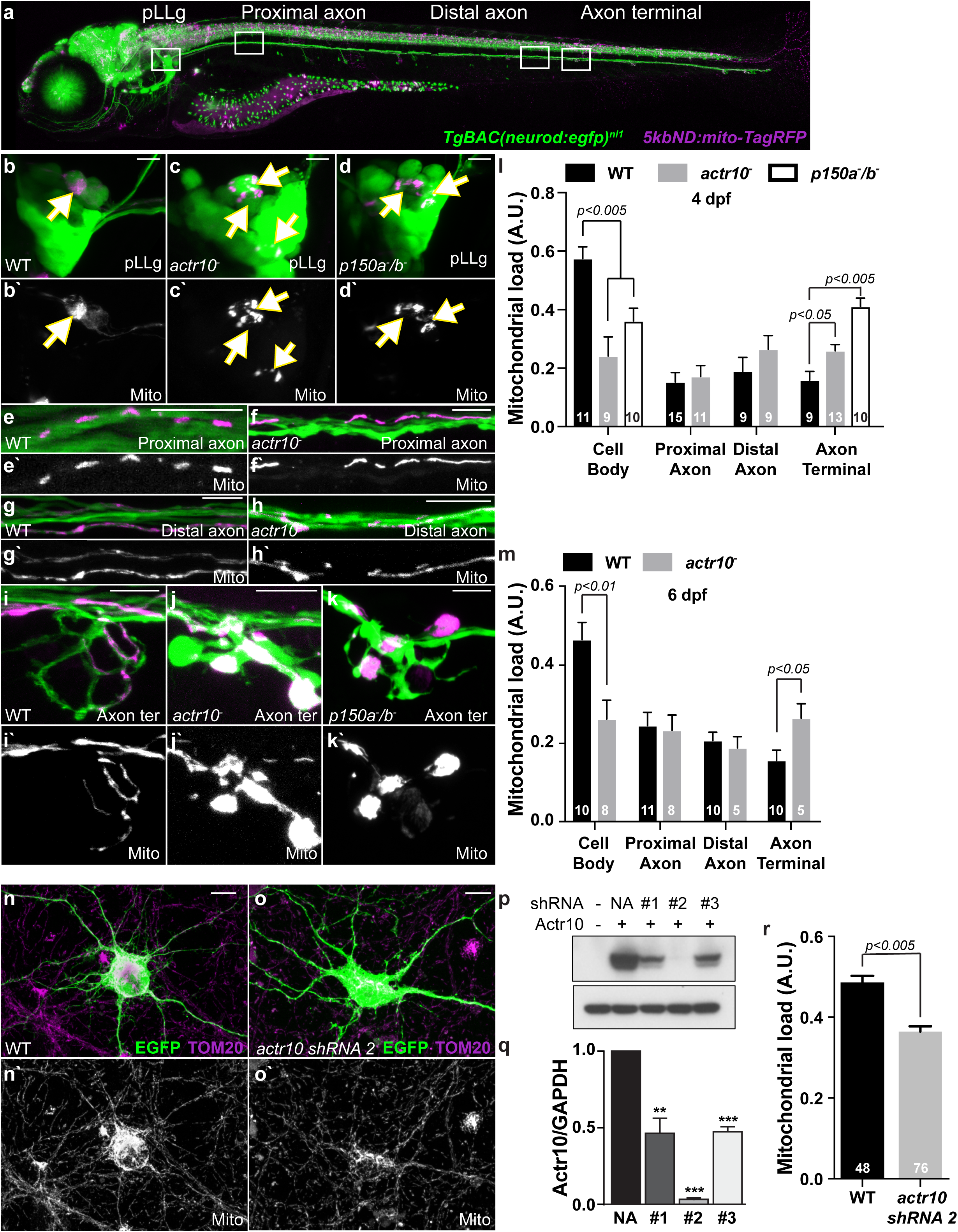
Disruption of retrograde mitochondrial movement impacts mitochondrial distribution in neurons. (**a**) Image of a 4 dpf *TgBAC(neurod:egfp)*^*nl1*^ transgenic zebrafish larva expressing mitochondrially localized TagRFP (shown in magenta) mosaically in neurons. Regions imaged in **b**-**k** shown in white boxes. (**b**-**d**) pLLg neurons have cytosolic GFP and a subset express TagRFP in mitochondria (arrows). In wildtype animals, mitochondria (magenta in **b**; white in **b**’) fill the soma. (**c**) In *actr10*^*-*^ mutants, cell body mitochondrial area is reduced. (**d**) Similarly, *p150* mutants show reduced cell body mitochondrial load. (**e**-**h**) Mitochondrial density is not significantly different in axonal segments with retrograde mitochondrial transport disrupted. (**i**-**k**) Mitochondria accumulate in axon terminals of mutant lines with failed retrograde mitochondrial transport. (**l**,**m**) Quantification of mitochondrial load at 4 dpf (ANOVA; Tukey HSD posthoc contrasts) and 6 dpf (ANOVA). (**n**-**r**) Knockdown of Actr10 in rat hippocampal neurons results in loss of cell body mitochondrial load. (**n**,**o**) Images of hippocampal neurons stained for TOM20 with a cytoplasmic EGFP fill and either vector only (**n**) or *actr10* shRNA#2 co-transfection (**o**). (**p**,**q**) Western blot and quantification showing knockdown of Actr10 by the shRNAs tested in HEK cells (**-*p*<0.01; ***-*p*<0.005; ANOVA; n=3). (**r**) Quantification of mitochondrial load in Actr10 knockdown hippocampal neurons (ANOVA). Sample sizes (**l**,**m** – number of larvae; **r** – number of neurons) indicated on graph. Scale bars – 10μm.

### Inhibition of retrograde mitochondrial motility leads to failed mitochondrial health

We next assayed various aspects of mitochondrial health with inhibition of retrograde mitochondrial transport. First, we analyzed chronic reactive oxygen species (ROS) production using the fluorescent ROS sensor TIMER in mitochondria. TIMER is a fluorescent protein that fluoresces in the green (488 nm) spectrum in its native state but irreversibly switches to red (568 nm) upon oxidation ^45–47^. This indicator has been used previously to assay chronic exposure to reactive oxygen species in zebrafish ^36,48^. We used mosaic expression of mitochondrially localized TIMER in individual neurons of the pLLg (Fig. 4a-e) and assayed the red:green fluorescence ratio at 4 and 6 dpf in *actr10*^*-*^ mitochondrial transport mutants and wildtype siblings. This analysis revealed there is a consistent increase in the red:green TIMER levels in axon terminals when retrograde mitochondrial transport is inhibited, indicating that these organelles accumulate more ROS. Interestingly, at 6 dpf, the red:green TIMER ratio is significantly lower in retrograde transport mutants in the neuronal cell body, potentially revealing the cell body as the primary site for mitochondria biogenesis (Fig. 4f).

**Figure 4:**
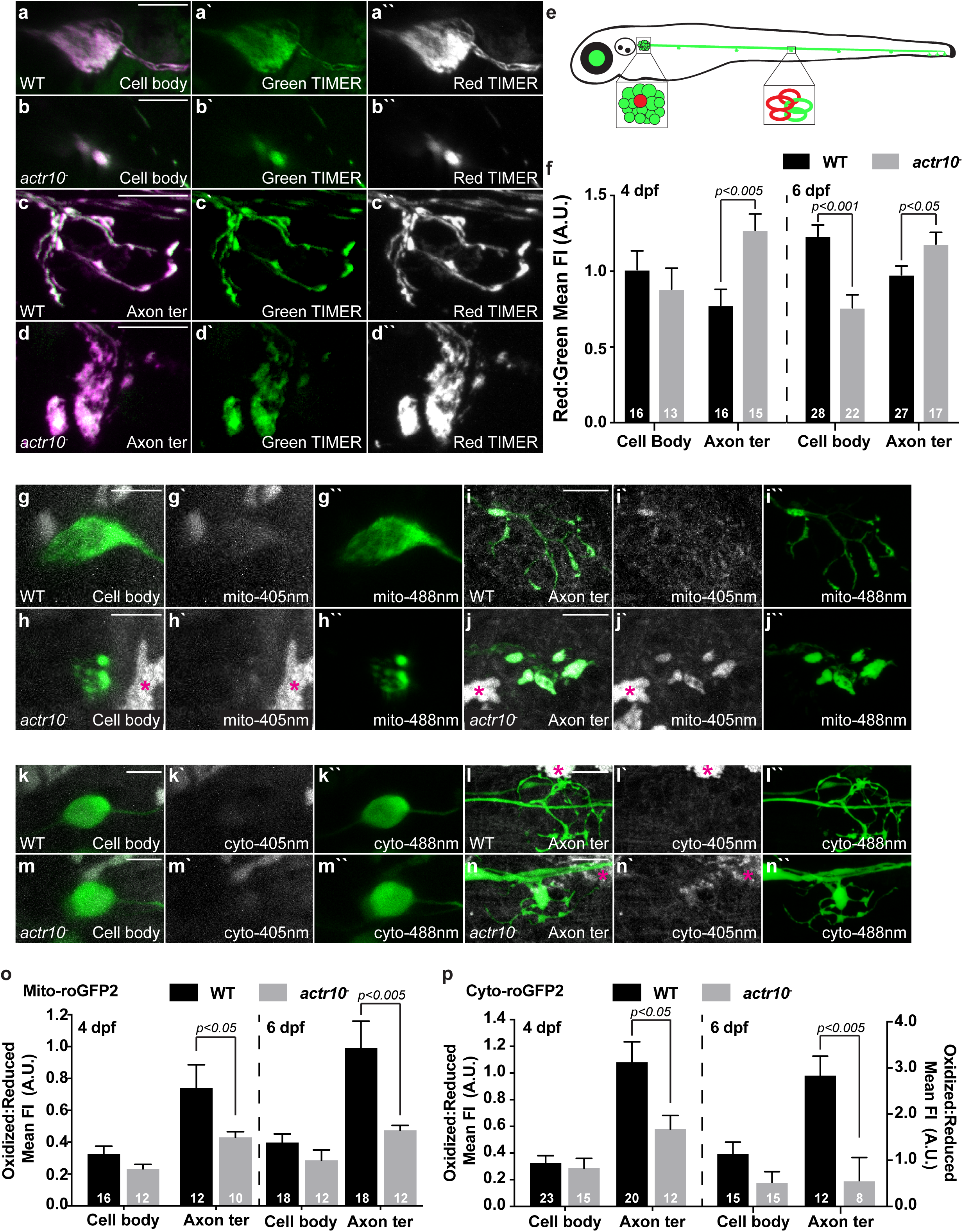
Loss of retrograde mitochondrial transport results in altered measures of acute and chronic reactive oxygen species (ROS). (**a**,**b**) Expression of mitochondrially localized TIMER in a wildtype (**a**) and *actr10*^*-*^ mutant (**b**) pLL neuron cell body. (**c**,**d**) TIMER fluorescence in axon terminal mitochondria of wildtype and *actr10*^*-*^ mutants at 4 dpf. Oxidized TIMER is shown in magenta or white, reduced TIMER in green. (**e**) Schematic of the imaging setup, illustrating the single pLL cell body and single axon terminal expressing TIMER (shown in red) in mitochondria. (**f**) Quantification of the red (oxidized) to green (reduced) TIMER protein at 4 and 6 dpf demonstrates cumulative oxidative stress in axon terminal mitochondria with retrograde transport reduction (ANOVA). Cell body mitochondria show decreased ROS at 6 dpf (ANOVA). (**g**-**p**) roGFP2 reveals acute changes in ROS levels in mitochondria of *actr10*^*-*^ mutants. (**g**-**j**) Mitochondrially localized roGFP2 in pLLg cell bodies (**g**,**h**) and axon terminals (**i**,**j**) of wildtype and *actr10*^*-*^ mutants at 4 dpf. (**k**-**n**) Cytoplasmic roGFP2 expression in pLLg neuron cell bodies (**k**,**m**) and axon terminals (**l**,**n**) of wildtype and *actr10*^*-*^ mutants at 4 dpf. (**o**,**p**) Quantification of the ratio of oxidized to reduced roGFP2 in neuronal compartments at 4 and 6 dpf (ANOVA or Wilcoxon Rank Sum). Asterisks on autofluorescent pigment cells. Sample sizes indicated on graph. Scale bar – 10μm.

Next, we assayed acute ROS levels using both cytosolic (cyto-) and mitochondrially (mito-) localized roGFP2 ^49,50^. This reversible indicator has peak fluorescence in the 405 nm spectrum when oxidized and 488 nm spectrum when reduced. We used transient transgenesis to express either cyto- or mito-roGFP2 in pLL neurons and assayed the oxidized to reduced roGFP2 ratio in the cell bodies and axon terminals (Fig. 4g-n). At both 4 and 6 dpf we observed similar trends: Mitochondria in retrograde transport mutants show a lower oxidized:reduced mito-roGFP2 fluorescence ratio in axon terminals compared to wildtype siblings (Fig. 4o). Mito-roGFP2 changes are mirrored by cyto-roGFP2 fluorescence ratios (Fig. 4p). Together, our data show that mitochondria retained in axon terminals accumulate higher levels of ROS over extended durations but show lower ROS production acutely. As ROS is a natural product of oxidative phosphorylation, we next sought to address the potential of the mitochondrial matrix and consequent ATP production after disruption of retrograde mitochondrial transport.

To assay mitochondrial matrix potential, we used the vital dye TMRE (tetramethylrhodamine ethyl ester). This cationic dye accumulates in the negatively charged mitochondrial matrix to a degree relative to potential ^51,52^. Larvae carrying the *TgBAC(neurod:egfp)*^*nl1*^ transgene to label neurons with cytoplasmic GFP were incubated in 25μm TMRE for an hour and then imaged live. This analysis revealed that at both 4 and 6 dpf, mitochondrial matrix potential was compromised in the pLLg and axon terminals when mitochondrial retrograde transport is inhibited in the *actr10*^*-*^ mutant (Fig. 5a-f). Decreased acute ROS production and matrix potential is predicted to effect oxidative phosphorylation and ATP production. To assay mitochondrial productivity, we expressed the ratiometric sensor PercevalHR in pLL neurons and assayed the ATP (488 nm) to ADP (405 nm) fluorescence ratios in the cell body and axon terminal (Fig. 5g-k) ^36,53^. Somewhat surprisingly, in *actr10*^*-*^ mutants we observed no change in the overall ATP:ADP ratio in either compartment at 4 or 6 dpf (Fig. 5l). We also assayed motor neuron axon ATP:ADP ratios using PercevalHR. Similar to pLL axons, we observed no change in the overall ATP:ADP ratio in *actr10*^*-*^ mutant motor neurons (Fig. 5m and Supplementary Fig. S2).

**Figure 5:**
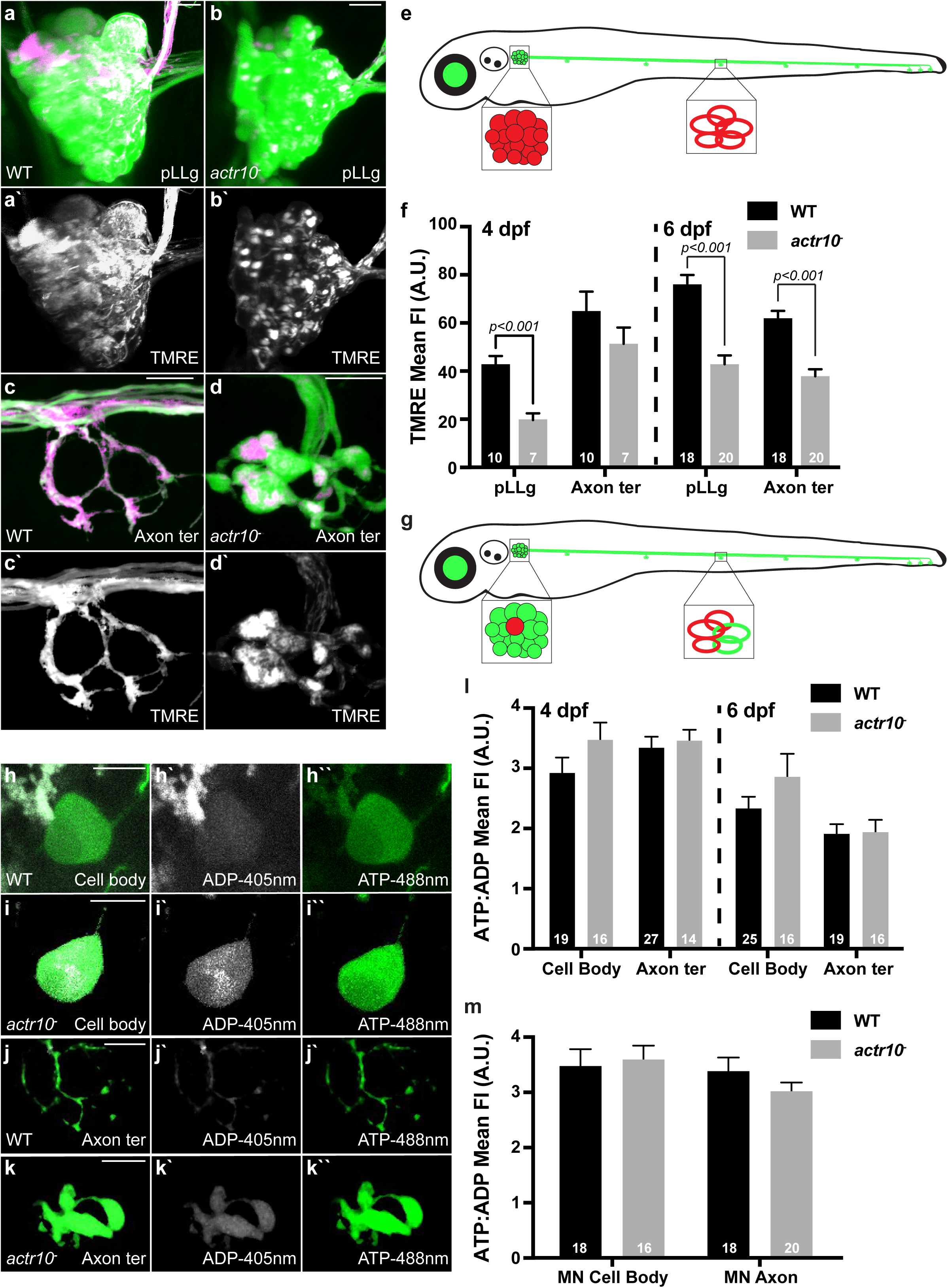
Mitochondrial matrix potential is lowered in retrograde transport mutants without effecting cytosolic ATP:ADP ratios. (**a**-**f**) TMRE staining (magenta in merge; white in single channel) of the mitochondrial matrix in pLLg neuronal cell bodies and their associated axon terminals at 4 dpf. Neurons are labeled with cytosolic GFP. TMRE signal in neurons was isolated using the ImageJ *Image calculator* tool. (**e**) Schematic of the regions of the larva used for this analysis. (**f**) Mean TMRE fluorescence is decreased in cell body mitochondria at 4 dpf and in the cell body and axon terminal mitochondria at 6 dpf when retrograde mitochondrial transport is inhibited (ANOVA). (**g**) Schematic of mosaic PercevalHR expression in the neuronal soma and axon terminal. (**h**-**k**) Wildtype and *actr10*^*-*^ mutant pLLg neuronal cell bodies (**h**,**i**) and axon terminals (**j**,**k**) expressing PercevalHR at 4 dpf. (**l**) Quantification of the ATP:ADP ratio at 4 and 6 dpf shows no deficits in *actr10*^*-*^ mutants (ANOVA). (**m**) Similarly, ATP:ADP ratios are unaffected in motor neuron axons at 4 dpf in *actr10*^*-*^ mutants (ANOVA). Sample sizes indicated on graphs. Scale bar – 10μm.

Because mitochondrial load is increased in *actr10*^*-*^ mutants axon terminals, we next asked if the per mitochondrial ATP:ADP ratio was altered. When mitochondrial load (mito area:axon terminal area) is considered, there is a significant reduction in the relative amounts of ATP to ADP (WT: 20.26±1.00; *actr10*^*-*^: 13.39±1.39; Wilcoxon Rank Sum *p*<0.0005), indicating decreased per mitochondria productivity. Finally, to confirm our Perceval HR results, we assessed ATP levels using a non-ratiometric sensor, ATP SnFR ^54^. As we observed with PercevalHR, there was no change in the overall level of ATP in cell bodies or axon terminals of pLL sensory neurons in *actr10*^*-*^ mutants compared to wildtype siblings (Supplementary Fig. S3) but a significant per mitochondria decrease was observed in axon terminals when mitochondrial load was controlled for (WT:4.8±0.20; *actr10*^*-*^: 2.93±0.36; Wilcoxon Rank Sum *p*<0.0001). Together, this indicates that though mitochondria are compromised in axon terminals and produce less ATP per organelle, neurons are able to compensate, perhaps through glycolysis, to maintain normal cytosolic ATP levels.

In addition to ATP production, mitochondria have many other roles in the neuron. One function that is intimately related to neural circuit function is calcium buffering ^3,9,10,12,36^. Therefore, we assayed whether retrograde mitochondrial motility is important for mitochondrial calcium buffering using a dual transgenic strategy. We combined two transgenic lines to express either G-GECO, a green calcium indicator, in the cytosol (*Tg(5kbneurod:G-GECO)*^*nl19*^) or R-GECO, a red calcium indicator, in mitochondria (*Tg(5kbneurod:mito-R-GECO)*^*nl20* 36,55^. We crossed these transgenes into the *actr10*^*-*^ mutant line and assayed the mitochondrial to cytoplasmic baseline calcium levels in the pLLg and axon terminals at 4 and 6 dpf (Fig. 6a-e). This demonstrated that, compared to controls, there was a reduction in mitochondrial:cytosolic calcium levels in mutant axon terminals (Fig. 6f). Together, these analyses of mitochondrial health and function indicate that mitochondria which cannot move in the retrograde direction have chronic increases in ROS exposure, lowered matrix potential, lower per mitochondria ATP production, and a compromised ability to buffer cytosolic calcium.

**Figure 6:**
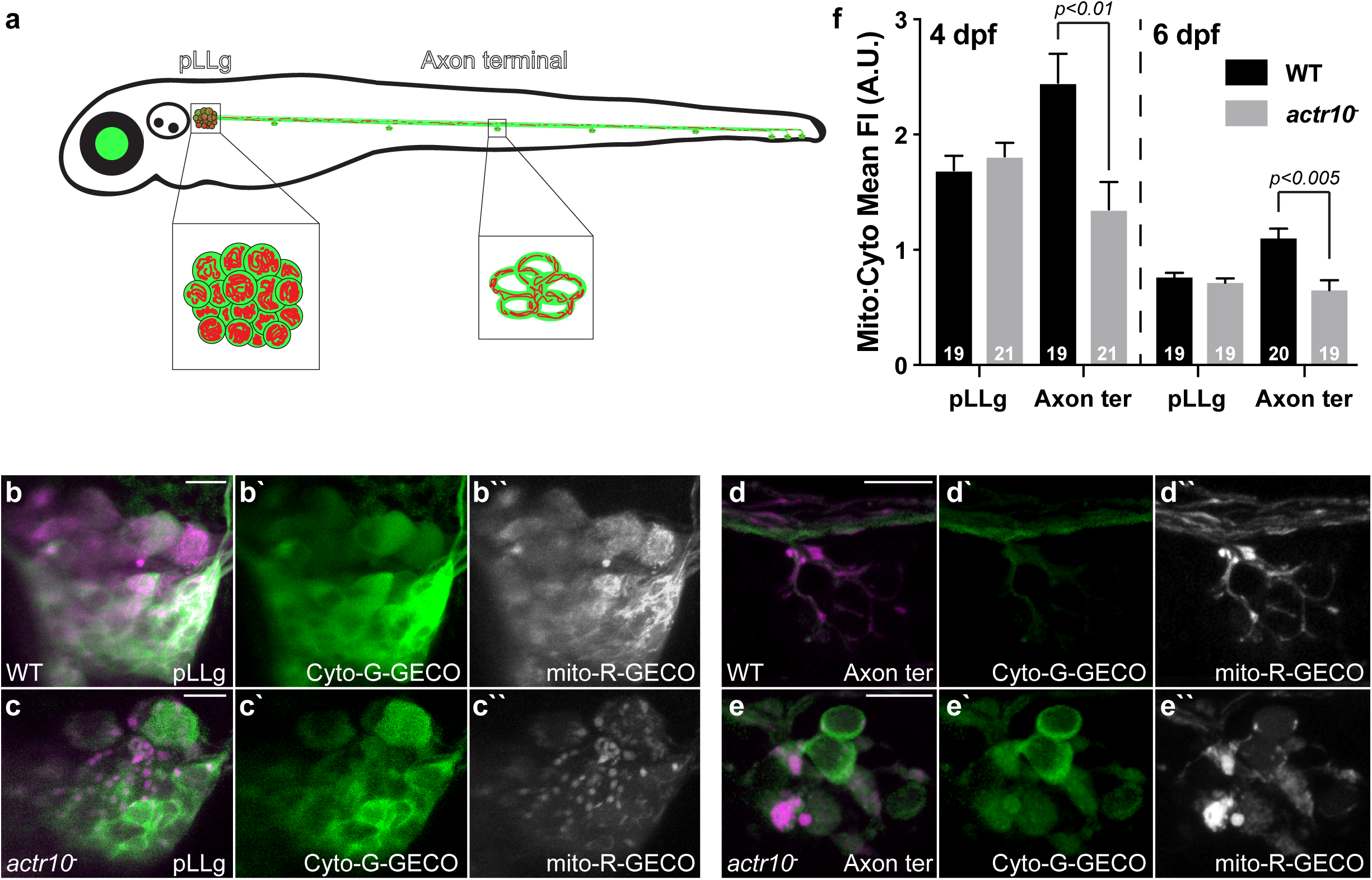
Mitochondrial calcium buffering is decreased in axon terminal mitochondria when retrograde transport is disrupted. (**a**) Schematic of the double transgenic used to analyze mitochondrial:cytoplasmic calcium levels. (**b**-**e**) Neuronal cell bodies and axon terminals expressing mitochondrially localized R-GECO (magenta in merge, white in single channel) and cytoplasmic G-GECO (green) in wildtype and *actr10*^*-*^ mutants at 4 dpf. (**f**) Quantification of mitochondrial:cytoplasmic GECO signal at 4 and 6 dpf (ANOVA or Wilcoxon Rank Sum). Sample sizes indicated on graph. Scale bars – 10μm.

### Abnormal circuit function with inhibition of mitochondrial retrograde transport

The alterations in mitochondrial function, particularly the decrease in calcium buffering, lead us to next ask if the function of the neural circuit was altered when retrograde mitochondrial transport was inhibited. First, we immunolabeled neuromasts to label sensory hair cells (HCs, Myosin 7a), along with pre-(Ribeye-positive) and post-(MAGUK-positive) synaptic puncta in *actr10*^*-*^ mutants and wildtype siblings (Fig. 7a-c). Somewhat unexpectedly, this showed that there are virtually no HCs in *actr10*^*-*^ mutants (Fig. 7b,c). We therefore rescued HCs by creating a transgenic strain that expresses mRFP tagged Actr10 in HCs (*Tg(myo6b:mRFP-actr10)*^*y610*^). *actr10*^*-*^ mutants expressing this rescue transgene (HC rescue) display normal numbers of sensory HCs and synapses per HC at 5 dpf (Fig. 7b-f); however, pre- and post-synapse size were slightly, though not significantly, reduced in *actr10*^*-*^ HC rescue larvae compared to wildtype siblings (Fig. 7g). For HC function, apical stereocilia must be mechanosensitive, meaning that upon stimulation mechanotransduction channels in stereocilia have to open to trigger basal calcium influx and activation of HC synapses. To assess mechanotransduction in HCs, we first applied FM1-43, a vital dye that permeates and labels HCs with functional mechanotransduction channels ^56,57^. This analysis demonstrated that the HCs in *actr10*^*-*^ mutants with HC rescue were able to take up FM1-43, indicating they are able to mechanotransduce (Fig. 7h).

**Figure 7:**
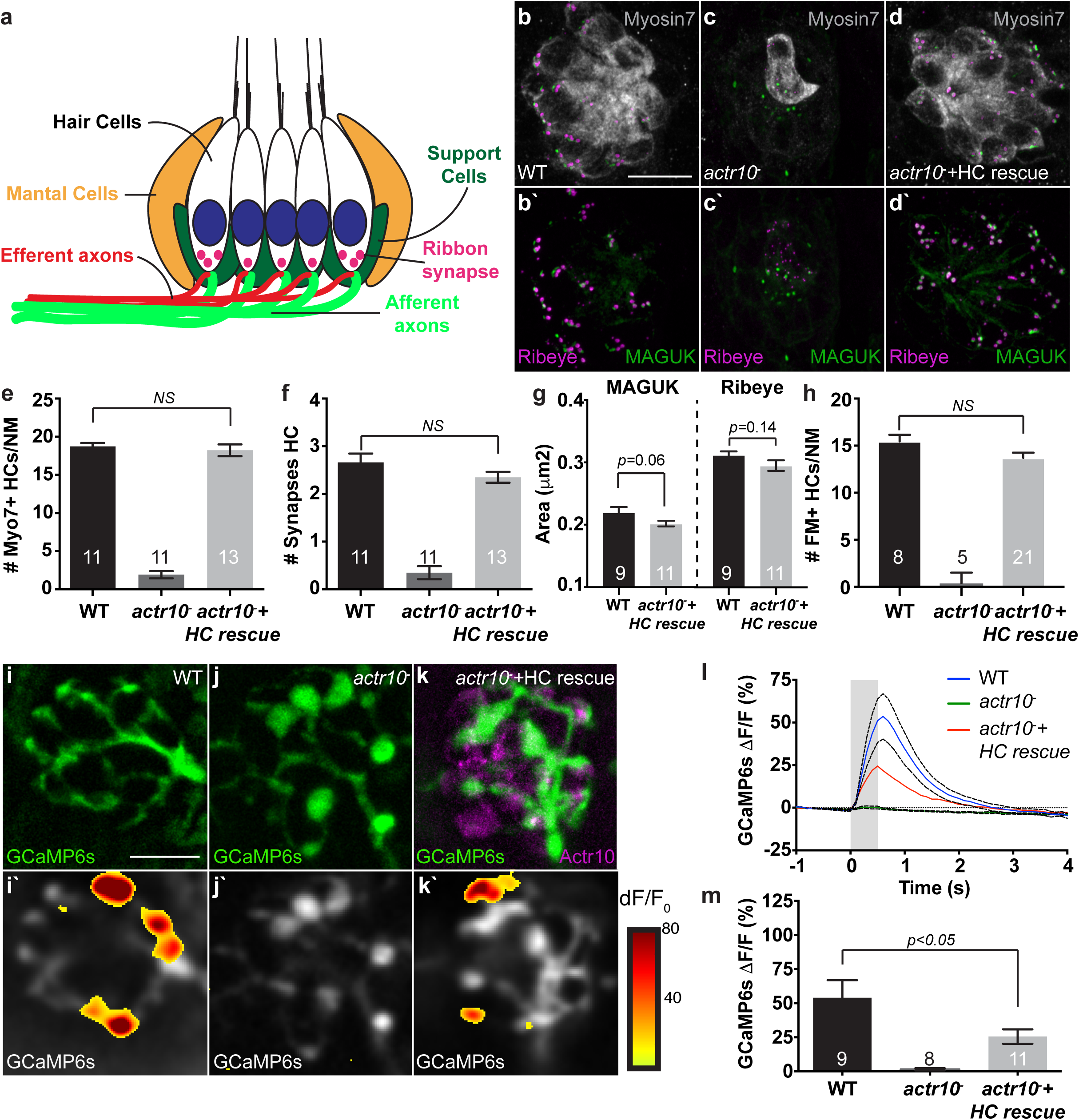
Axonal response to stimulation is decreased in mitochondrial retrograde transport mutants. (**a**) Schematic of the neural circuit in a sensory neuromast of the pLL. HC stereocilia are deflected by water movement and signal through ribbon synapses to afferent axons. Support cells surround HC’s and efferent axons are also present. (**b**,**c**,**e**) *actr10*^*-*^ mutants have fewer sensory HC’s as assayed by Myosin 7a immunolabeling. (**b**-**e**) HC number is rescued by expressing RFP-tagged Actr10 in HCs (HC rescue) using the Tg(*Myo6b:mRFP-Actr10*) transgenic (ANOVA). (**b**-**d**,**f**) Synapse number, as assayed by Ribeye (pre-synaptic) and MAGUK (post-synaptic) immunolabeling, is rescued in the HC rescue *actr10*^*-*^ larvae (ANOVA). (**g**) While synapse number is rescued, pre- and post-synapse size is slightly reduced in *actr10*^*-*^ HC rescue larvae (*t*-test). (**h**) Mechanotransduction is rescued in *actr10*^*-*^ HC rescue mutants (ANOVA). (**i**-**m**) *actr10*^*-*^ mutant axon terminals show a reduced level of activity as assayed by changes in GCaMP6s fluorescence intensity with HC stimulation (*t*-test). (**i**’-**k**’) Heat map denotes changes in GCaMP6s fluorescence during stimulation. Sample sizes on graphs. Scale bar – 10μm.

We next asked if afferent axons were able to respond to HC stimulation using calcium imaging and previously published protocols ^57–59^. We used a fluid jet to mechanically stimulate HCs and monitored the change in GCaMP6s intensity in the afferent axon. Compared to wildtype siblings, *actr10*^*-*^ mutants with HC rescue showed depressed axonal responses during stimulation (Fig. 7i-m). As expected, *actr10*^*-*^ mutants which lack HCs showed no axonal response (Fig. 7j,l,m). The reduced axonal response observed in *actr10*^*-*^ HC rescue animals could be cell autonomous or due to a defect in another component of the circuit, such as the pre-synapse in the HC. We assayed HC synaptic function more closely using calcium imaging at the HC apex (site of mechanotransduction) and the HC base (the site of the presynapse; Supplementary Fig. S4a). During stimulation, we observed comparable changes in GCaMP6s fluorescence at the HC apex between *actr10*^*-*^ HC rescue mutants and wildtype siblings, confirming our FM1-43 data showing normal mechanotransduction (Supplementary Fig. S4b-e). Somewhat surprisingly, though mechanotransduction was normal, pre-synaptic activity at the base of the HC was depressed, even in HC rescued *actr10*^*-*^ mutants (Supplementary Fig. S4f-i). The depressed pre-(HC base) and post-(afferent axon) response could be due to the altered pre- and post-synapse sizes described above (Fig. 7g) as synapse size can dictate synaptic activity in this system ^60,61^ or an uncharacterized defect in another component of this circuit (Fig. 7a). Together, this data demonstrates that synaptic function is disrupted in the *actr10*^*-*^ mutants when retrograde mitochondrial transport is inhibited.

### Function of retrograde mitochondrial transport in axons

The defects we observed in mitochondrial health and function with inhibited retrograde transport in neurons, indicate that this process is critical for the maintenance of a healthy mitochondrial population. Additionally, our photoconversion analysis and mitochondrial load measures demonstrated that retrograde transport of this organelle throughout the neuron is necessary to maintain a homeostatic mitochondrial pool. The obvious next question is why do mitochondria need to constantly be in flux throughout this large cell. One potential reason for mitochondrial movement towards the cell body is to bring them closer to the site of protein synthesis for replenishment of mitochondrial proteins as almost 99% of mitochondrial proteins are encoded by nuclear DNA ^21,22^. Therefore, we wanted to determine if retrograde transport of mitochondria back to the cell body had a function in maintenance of the mitochondrial proteome. To do this, we fractionated mitochondria from *actr10*^*-*^ mutants and their wildtype siblings and analyzed the mitochondrial proteome using mass spectrometry (n=3 replicates; 100 larvae per sample). This identified a list of candidate proteins, some of which were decreased by more than 50% in *actr10*^*-*^ mutants. For proteins reduced by 50% or more, we compared the list manually to the MitoCarta2.0 database of validated mitochondrial proteins ^21^. This resulted in a list of 89 validated mitochondrial proteins reduced in *actr10*^*-*^ mutants. Gene Ontology analysis using the DAVID database ^62–64^ revealed that these proteins were functionally linked and able to be divided into eleven categories (Fig. 8a and Table 1). Together, these results indicate that inhibition of retrograde mitochondrial transport dramatically impacts the mitochondrial proteome.

**Table 1:**
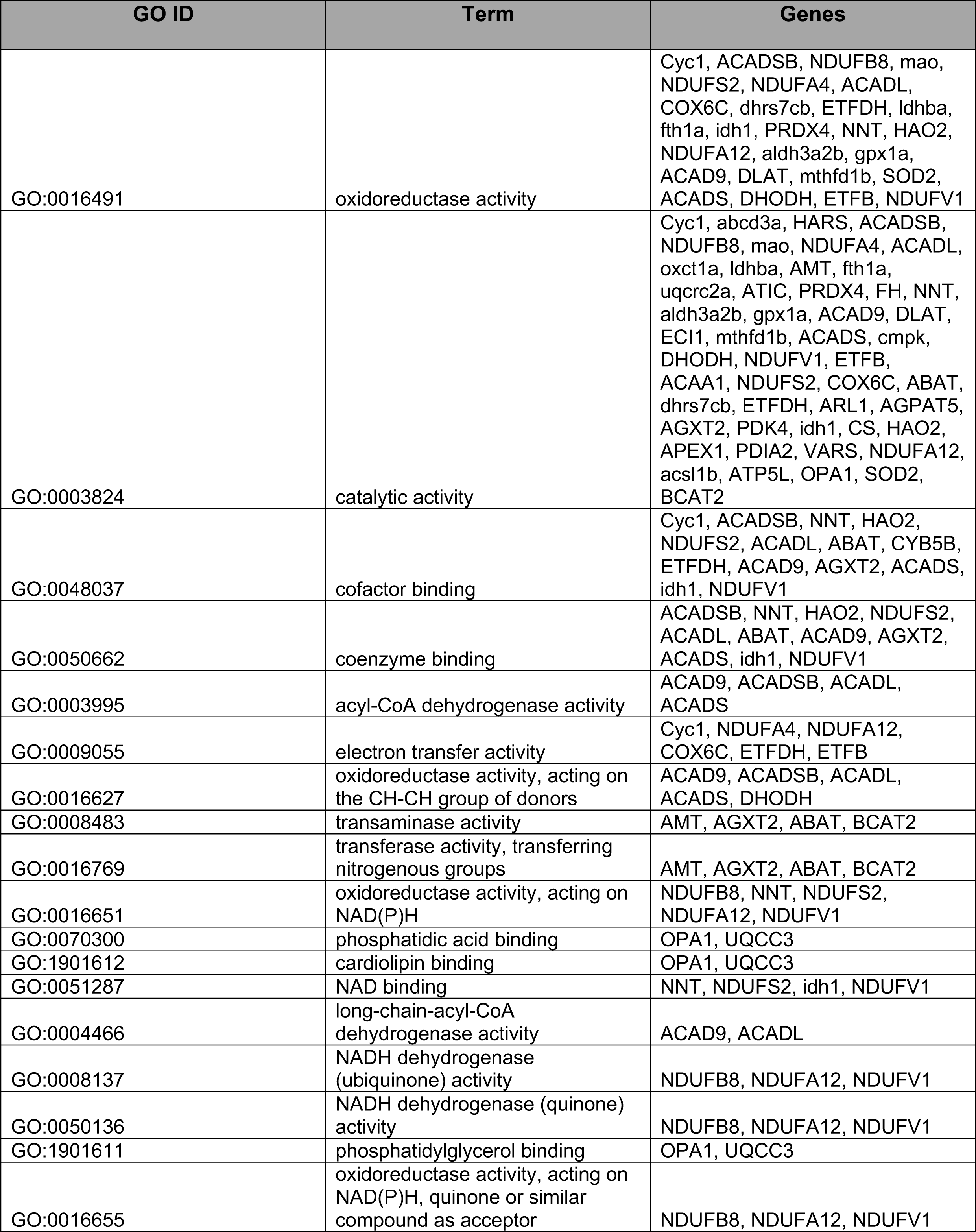
GO term functional analysis of proteins lost in *actr10*^*-*^ mutant mitochondria.

**Figure 8:**
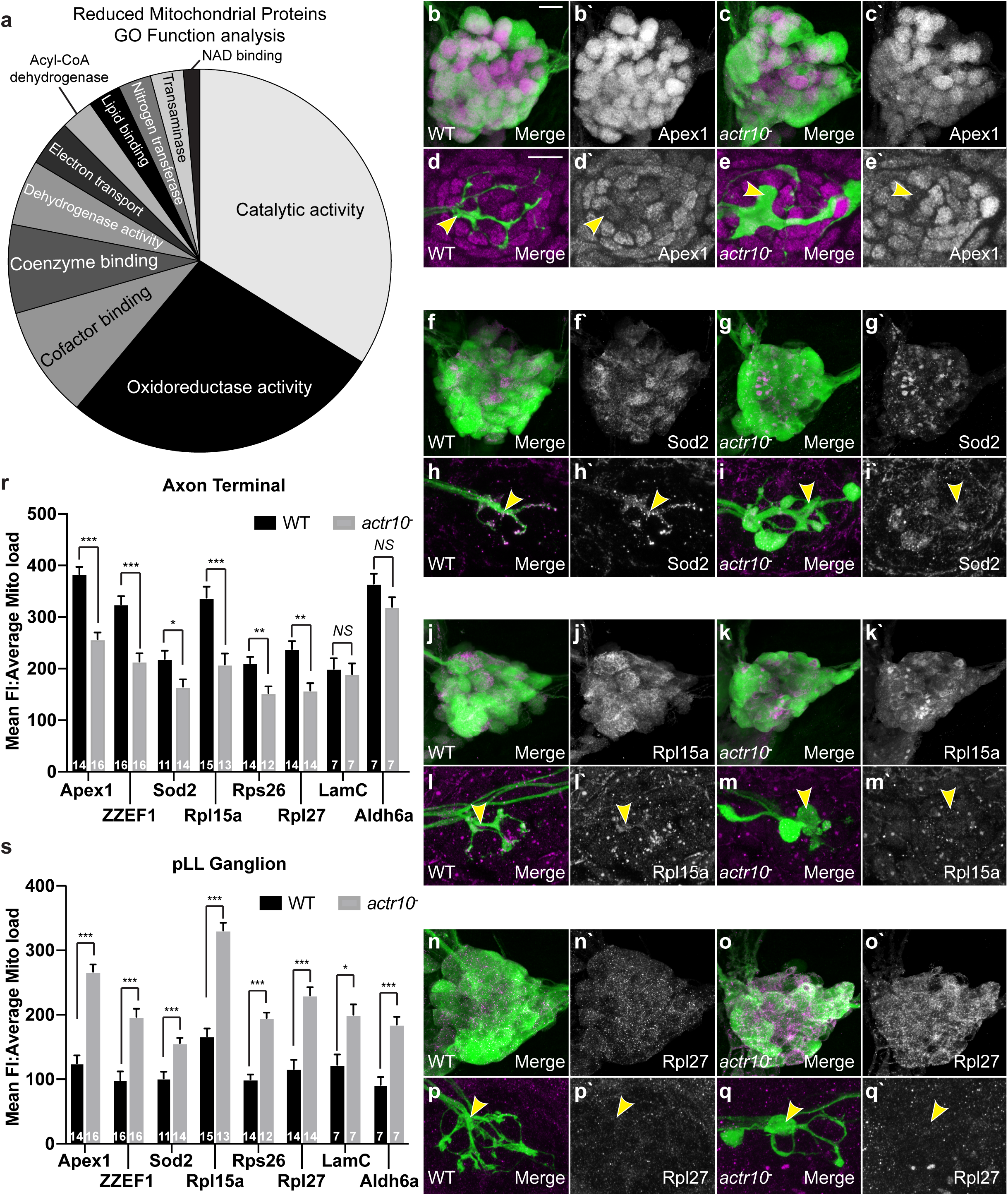
Defects in the mitochondrial proteome when retrograde mitochondrial transport is disrupted. (**a**) GO term analysis of functions associated with mitochondrial proteins decreased by >50% in *actr10*^*-*^ mutant mitochondria at 5 dpf. (**b**-**q**) Immunofluorescence analysis of a subset of the mitochondrial proteins downregulated in the mitochondrial fraction analyzed by mass spectrometry in the pLLg and axon terminal at 5 dpf. Immunofluorescence signal in the pLLg was isolated from surrounding tissue using the ImageJ *Image calculator* tool. Arrowheads point to a region of the axon terminal for each stain. (**r**,**s**) Quantification of fluorescence intensity of the immunolabeling in the pLLg and axon terminals normalized to average mitochondrial load (ANOVA; ***-*p*<0.0005; **-*p*<0.005; *-*p*<0.05). Sample size indicated on graph. Scale bar – 10μm.

Because the mitochondrial proteomes analyzed were generated from whole larval mitochondrial fractions, we next wanted to validate this data in neurons. If retrograde transport is indeed necessary for these proteins to be loaded into mitochondria, we predicted we would observe a decrease in the amount of these proteins in axon terminal mitochondria of *actr10*^*-*^ mutants. We performed immunolabeling for a subset of candidates including DNA repair enzymes (Apex1, ZZEF1), antioxidants (Sod2), and mitochondrial ribosomal proteins (Rpl15a, Rpl27, and Rps26) and analyzed their mitochondrial load using immunofluorescence. Strikingly, these proteins were elevated in neuronal cell bodies but lost in axon terminal mitochondria of the *actr10*^*-*^ mutant, implicating retrograde mitochondrial transport in their import into this organelle (Fig. 8b-s; Supplementary Fig. S5a-h). Two proteins not altered in our mass spec experiment, LamC1 and Aldh6a, were not elevated in our immunofluorescence analysis in axon terminals and served as negative controls (Fig. 8r; Supplementary Fig. S5i-p).

Altogether, our work demonstrates that retrograde mitochondrial transport occurs in non-pathological conditions to maintain a homeostatic mitochondrial population in the neuron.

## Discussion

The necessity of mitochondrial motility in neuron is demonstrated by the wealth of disease literature showing a correlation between abnormal mitochondrial localization and neurodegenerative disease. For anterograde transport, the reasons for pathology are intuitive since anterograde mitochondrial transport is essential for moving this organelle into the axon from the primary site of mitochondrial biogenesis near the nucleus ^65^. The purpose of retrograde mitochondrial movement has been postulated to remove damaged organelles from the axon. This conclusion is based on pharmacological manipulations that perturb mitochondrial health and lead to enhanced retrograde mitochondrial transport. For example, treatment of cultured neurons with drugs such as rotenone, Antimycin A, FCCP, and others cause a loss of mitochondrial matrix potential and can lead to enhanced retrograde mitochondrial motility ^15,16,31^. Additionally, analyses of mitochondrial movement in neurons treated with a matrix potential indicator have shown that lower matrix potentials correlate with retrograde movement ^31^. However, this is controversial as conflicting studies have shown no correlation between retrograde movement and mitochondrial health using similar methodologies ^15,34^. More than likely, the truth lies somewhere in the middle with mitochondrial health feeding into a larger signaling network that regulates mitochondrial localization. Our work supports a role for consistent mitochondrial movement over the scale of hours to maintain a homeostatic distribution of this organelle throughout the neuron’s volume.

By imaging neuronal mitochondrial dynamics over time scales of hours and days, we have shown that retrograde mitochondrial motility occurs in axons at a reliable rate. Specifically, in synaptically active axon terminals of wildtype animals, we observed a complete turnover of the mitochondrial population in less than 24 hrs. This entirely intact, in vivo system, has not been treated with any pharmacology to manipulate mitochondrial health, indicating that retrograde mitochondrial motility occurs in the absence of mitochondrial pathology. Additionally, we have shown that this turnover occurs via retrograde transport and is essential for maintaining a homeostatic distribution of mitochondria throughout mammalian and zebrafish neurons. Inhibition of retrograde mitochondrial movement leads to an accumulation of this organelle in axon terminals and a loss of mitochondrial load in the neuronal cell body. From this, we argue that retrograde mitochondrial transport is not just a disposal mechanism for the removal and ultimate degradation of this organelle from axons. Rather, balanced anterograde and retrograde mitochondrial transport is critical for maintaining the homeostatic flux of this organelle observed in neurons. It is the disruption of this balanced flow that leads to the accumulation of damaged organelles in axon terminals.

The constant movement of mitochondria in neurons is highly energetically demanding, leading to questions as to why this would be advantageous to the cell. Our data supports the idea that at least in part this retrograde movement is to allow replenishment of the mitochondrial proteome. Mitochondria have their own DNA that encodes 13 proteins in humans, primarily components of the electron transport chain ^66^. This is a small fraction of total mitochondrial proteins, however, as there are estimated to be more than 1100 proteins in these organelles ^21,22,67^. These proteins have diverse half-lives with some being on the order of hours while other proteins are stable for days to weeks ^23,24^. Once damaged or otherwise utilized, proteins are degraded and need to be replaced. The majority of mitochondrial proteins are derived from genes encoded by nuclear DNA located in the cell body. The cell body is also the site of the majority of protein synthesis in neurons. While mRNA and protein transport from the cell body to organelles in the axon can contribute to the replenishment of mitochondrial proteins ^13^, it stands to reason that moving the organelle back to the primary site of protein synthesis is an efficient mechanism for bulk mitochondrial protein turnover. Our proteomics data support this as mitochondria isolated from mutants that lack mitochondrial retrograde transport show a substantial reduction in approximately a third of their mitochondrial proteins. While we cannot rule out that these defects are correlational rather than causal, the elevation of cell body levels of proteins lost from mitochondria in axon terminals is strong evidence that at least a subset of protein turnover in axon terminal mitochondria requires retrograde transport to the cell body.

The impact of mitochondrial retrograde transport and proteome support on neural circuit function is clear and perhaps not surprising. Mitochondria have been shown to be essential for local ATP synthesis and calcium buffering to support active synapse ^11,12^. In our studies, mutants with retrograde mitochondrial transport disruption in axons show diminished synaptic activity as assayed by calcium imaging in the pre- and post-synapse. Rescue by expression of wildtype Actr10 in the pre-synaptic HC using a transgenic approach resulted in full rescue of HC morphology as well as rescue of calcium influx at the site of mechanotransduction but is unable to completely rescue synapse morphology or function. Although synapse number is normal, the pre- and post-synapse are slightly smaller is size. As synapse size has been shown to correlate with activity ^60,61^ this could indicate that the reduced pre- and post-synaptic response is due to altered size of the assembled synapses. Whether this change in synapse size is due to incomplete rescue in the HC or solely due to defects in the afferent axon is unclear; however, given the strong rescue of HC morphology, synapse number, and apical mechanotransduction, it is likely that the reduced synapse size is a result of defects in the afferent axon that is innervating the HC. Together, our work illustrates the importance of mitochondrial localization and health on the formation and function of an in vivo neural circuit.

## Conclusion

We have shown that retrograde mitochondrial motility is commonly observed in non-pathological conditions in axons and this movement causes a complete turnover of the axon terminal population within a day. Inhibition of this movement leads to a severe reduction in the mitochondrial proteome, specifically in proteins known to be important for mitochondrial DNA maintenance, mitochondrial protein translation, and the reduction of oxides that are detrimental to these organelles. Consequently, mitochondria isolated in axon terminals in retrograde transport mutants display evidence of chronic exposure to reactive oxygen species and reduced health and function. This data supports the necessary bidirectional transport of mitochondria in neurons to maintain the health of this integral organelle and consequently the health and function of the neural circuit.

## Supporting information

Supplemental Figures and resources

## Acknowledgements

We would like to thank Dr. A. Nechiporuk for his thoughtful comments on this work and contributing to the production of our zebrafish strains. We would also like to acknowledge members of the Drerup lab (D. Kawano, E. Rosenfeld-Gur, and S. Wisner) for their critique of this work. TIMER and roGFP2 constructs were provided by Addgene. A subset of the antibodies used were provided by the Developmental Studies Hybridoma Bank. Funding for this work was provided by the Intramural Research Program Grants from the NICHD(1ZIAHD008964-02 to C.M.D.), NIDCD (1ZIADC000085-01 to K.S.K), and NINDS (1ZIANS002994-17).

## Author contributions

AM and CMD conceptualized the project; designed, performed, and analyzed experiments; and wrote the manuscript. KK designed, performed and analyzed synapse formation and activity experiments and edited the manuscript. KP, AB, and NM performed and analyzed experiments. SW and RL designed rat Actr10 shRNAs and transfected and stained rat hippocampal neurons.

## Materials and Methods

### Zebrafish strains and husbandry

All zebrafish (*Danio rerio*) work was done in accordance with the NICHD/NINDS IACUC guidelines (protocol ID Drerup18.008 or protocol Kindt #1362-13). Adult animals were kept at 28°C and spawned according to established protocols ^68^. Embryos and larvae were kept in embryo media at 28°C and developmentally staged using established methods ^69^. The *Tg(myo6b:mRFP-actr10)*^*y610*^ stable transgenic line was derived using Tol2-mediated transgenesis and the Gateway system according to established protocols ^70^.

### Transient transgenesis

For analyses of mitochondrial localization and measures in single neurons, transient transgenesis was utilized. Plasmid DNA encoding the *5kbneurod* (pLL neurons; ^71^) or *mnx1* (motor neurons; ^72,73^) promotors driving constructs of interest were derived using Gateway technology ^70^. For expression, 3-13pg of plasmid DNA was microinjected into zebrafish zygotes as previously described ^37,74^. For analysis, larvae expressing the construct of interest in a subset of pLL or motor neuron cell bodies were selected using a Zeiss AxioZoom fluorescent dissecting scope. Larvae were anesthetized in 0.02% tricaine and mounted individually in 1.5% low melt agarose in embryo media and imaged with a Zeiss LSM800 confocal microscope with a 63X water immersion objective, NA1.2.

### Genotyping and generation of the p150a/b double mutant line

*actr10*^*nl15*^ and *p150b*^*nl16*^ mutants were genotyped as previously described ^43^. We generated new p150a alleles using CRISPR-Cas9 technology ^75–79^. A gRNA (GGTAAGATGAGTTCAGACGG; 125pg) targeted to exon 2 of the *p150a* locus was co-injected with Cas9 protein (500pg; Integrated DNA Technologies). Embryos were assessed for cutting efficiency by lysing a subset of injected embryos and genotyping as described ^79^ using PCR-based methods. Siblings were raised and F0s tested for changes in the *p150a* locus by outcrossing and genotyping resulting embryos. F0s confirmed to carry detrimental changes in *p150a* were subsequently mated to generate F1 larvae for analysis. All larvae were sequenced and only those confirmed to have detrimental mutations, e.g. frame-shifts or loss of the start codon, were analyzed. Additionally, *p150a* mutants have a characteristic eye phenotype ^80^ which was confirmed in all analyzed larvae for Figure 3.

### Cortical neuron culture analysis

Primary hippocampal neurons were isolated from E17-E18 rat embryos as described previously ^81^. Neurons were co-transfected with 500 ng each of EGFP and pSuper vector or Actr10 shRNA #2 (ggtcctggattagtggatatag) at DIV 13 using Lipofectamine 2000. 4 days post-transfection, at DIV 17, the cells were fixed and stained for endogenous TOM20, a mitochondrial marker. Fixation was performed in PBS containing 4%PFA and 4% sucrose for 7-8 min, followed by permeabilization with 0.25% Tx-100 in PBS for 10 min. TOM20 staining was done using anti-Tom20 for 1 hr at RT, followed by staining with anti-Rabbit Alexa-555 (1:1000, Life Technologies) for 1 hr at RT. The slides were mounted in Prolong gold antifade and imaged using a Zeiss LSM800 confocal microscope with a 63X/NA 1.4 objective. Cell somas in 7 to 21 neurons were analyzed per coverslip. Five slides with 2 coverslips each were analyzed per condition in two blinded, experimental replicates.

### Mitochondrial photoconversion and time-lapse imaging

Mitochondria in axons, axon terminals, or cell bodies (separate experiments) were converted using a 405nm laser on the Zeiss LSM800 confocal microscope with a 40X/NA1.0 dipping lens. For conversion, larvae were mounted in 0.8% low melt agarose and immersed in embryo media with < 0.02% tricaine. A pre-conversion image was taken and then a region of interest (ROI) was defined in *Zen* around the region to be converted. A *z*-stack with a range of 9μm to 25μm was set up and the ROI was scanned with 5-7% 405nm laser power 1-2X until complete conversion of the ROI was achieved. Conversion was confirmed with a post-conversion image. For the 24 hr time-point, embryos were either housed in agarose or removed from the agarose and housed in embryo media at 28°C overnight in the dark before reimaging the following day. For time-lapse imaging, *z*- stacks through the ROI were set up to be scanned every 1, 3, or 10 minutes for 1 to 16 hours. All imaging was done using the 488nm (1% laser power) and 568nm (1% laser power) lasers on a LSM800 confocal microscope with a 40X/NA1.0 dipping objective. For Figure 2, imaging was done using the 488nm (1% laser power) and 568nm (20% laser power). After imaging, larvae were genotyped.

### Mass Spectrometry of mitochondrial proteins

Mitochondrial fractions were isolated as previously described ^43,82^. After pelleting, mitochondria were lysed in 9M Urea buffer prior to label free Mass Spectrometry analysis. Proteins identified were compared to the Human MitoCarta database ^21^ to eliminate proteins not verified to be found in the organelle and fold changes calculated from averages of the three biological replicates. Gene Ontology term analysis of functional clustering was done using the DAVID online database (https://david.ncifcrf.gov/summary.jsp) with high stringency settings ^62–64^.

### Fluorescence intensity and mitochondrial area analyses

For analysis of fluorescence intensity in stable and transient transgenic zebrafish, imaging was done on a Zeiss LSM800 confocal microscope with either a 40X/NA1.0 dipping objective (photoconversion experiments), a 63XNA1.2 water immersion objective (live imaging), or a 63X/NA1.4 oil objective (immunofluorescence). Within experiments, all settings including laser power, gain, *z*-step size, and image resolution were kept consistent. Mean fluorescence intensity was then calculated from the neuronal area defined either manually or by masking the image using a neuron fill through the *z*-stack and the ImageJ image calculator function. All FI calculations were done using ImageJ ^83^.

For quantification of area, the cell body or axon terminal to be analyzed was defined by manual tracing or by masking the image using a neuronal cell fill in a projected *z*-stack. For quantification of mitochondrial area, a threshold was manually applied to a projected *z*-stack and the area masked selected and quantified.

### Immunohistochemistry, western blot, and vital dye labeling

For hair-cell synapse immunolabeling, zebrafish larvae at 5 dpf were fixed in 4 % PFA in PBS for 3.5 hrs at 4°C. After rinse, larvae were then permeabilized in acetone (stored at −20°C) for 5 minutes and blocked with PBST buffer containing 2 % goat serum, 2 % fish skin gelatin and 1 % BSA, overnight at 4°C. Primary antibodies were diluted in block solution. After removal of primary antibody, larvae were incubated in Alexa Fluor conjugated secondary antibodies. Larvae were then mounted on slides with ProLong Gold Antifade Reagent (Life Technologies). Fixed samples from hair-cell synapse immunolabeling were imaged on an inverted Zeiss LSM 780 laser-scanning confocal microscope using a 63X/1.4 NA oil objective lens. Images were acquired with a 5.0 x zoom at 620 x 620 every 0.19 µM. The Z-stacks were processed with Zeiss Zen Black software v2.1 using an Airyscan processing factor optimized with auto feature setting: Ribeye (7.5-7.8) and Maguk (6.0-6.3). Synapses and HCs were counted manually. Counts were done blinded. Myosin VIIa label was used to determine HC number.

To quantify HC pre- and post-synapse size, images were processed in ImageJ. In ImageJ, each Airyscan Z-stack was background subtracted using rolling-ball subtraction. Z-stacks containing the MAGUK channel were further bandpass filtered to remove details smaller than 6 px and larger than 20 px. A duplicate of the Z-stack was normalized for intensity. This duplicated Z-stack was used to identify individual ribbon and MAGUK using the Simple 3D Segmentation of ImageJ 3D Suite ^84^. Local intensity maxima, identified with 3D Fast Filter, and 3D watershed were used to separate close-by structures. The max Z-projection of the segmented Z-stack was used to generate a list of 2D objects as individual ROIs corresponding to each punctum. This step also included a minimum size filter, Ribeye: 0.08 μm^2^, MAGUK 0.04 μm^2^. Area and intensity measurements from these ROIs were exported from ImageJ.

For immunofluorescence for quantification of mitochondrial proteins, larvae at 4 dpf were fixed overnight at 4°C or 2 hrs at room temperature in 4% paraformaldehyde with 0.1% triton. Post-fixation, embryos were washed in water overnight, blocked in blocking solution (1% Bovine Serum Albumin, 0.1% Triton, 1% DMSO, 0.02% sodium azide, 5% goat serum), and incubated in primary antibody in blocking solution overnight at 4°C. Larvae were then washed and incubated overnight at 4°C with the appropriate AlexaFluor secondaries before washing and storage in 60% glycerol in 1X PBS/0.01% triton.

Western blot was performed as previously described ^74^. Larvae were lysed in lysis buffer (150mM NaCl, 20mM Tris-HCl pH 7.5, 1mM EDTA, 1% NP-40, 1% sodium deoxycholate) with phosphatase and protease inhibitors (25mM sodium fluoride, 10mM sodium orthovanadate, 1mM DTT, and 10uL protease inhibitor cocktail (Sigma Aldrich; P8340)). Extracts were incubated on ice for an hour, centrifuged for 30 minutes at 13200rpm at 4°C. Sample buffer was added to extracts which were then heated to 98°C for 15 minutes before running on a 10% SDS-PAGE gel. Following transfer to PVDF, membranes were blocked in 5% Omniblock before primary antibody in block was added for incubation overnight at 4°C. Membranes were washed 3 times for 5 minutes in PBS, 0.1% tween-20 and secondary antibody in PBS, 0.1% tween-20 was incubated with the membrane for 1.5 hours. After washing 3 times for 5 minutes in PBS, 0.1% tween-20, membranes were incubated with Supersignal West Pico Chemiluminiscent substrate and then exposed to film.

Vital dye labeling with TMRE (tetramethylrhodamine ethyl ester) was performed as previously described ^36^. Briefly, zebrafish larvae at 4 dpf in 25μM TMRE in embryo media with 0.1% DMSO (dimethyl sulfoxide) for an hour in the dark. Larvae were subsequently washed three times in embryo media prior to being anesthetized in 0.02% tricaine in embryo media, mounted in 1.5% low melt agarose, and imaged with a 63X, NA1.4 objective on a confocal microscope (Zeiss LSM800). For analysis, mitochondrial TMRE was measured in axon terminals and the pLL ganglion after subtraction of nonneural tissue using the ImageJ *Image Calculator* function using the GFP fill from the *TgBAC(neurod:egfp)*^*nl1*^ transgenic line.

Vital dye labeling of mechanotransducing HCs using FM1-43 was done as previously described ^56,57^. Larvae were incubated in 3μm FM1-43^647^ (Invitrogen) in embryo media with 0.1% DMSO for 30 seconds and immediately washed 3 times in embryo media before being anesthetized and mounted in 1.5% low melt agarose for imaging as described above for TMRE labeling. For analysis, the number of FM1-43 positive HCs in the neuromast were counted for NM3 in the mid-trunk.

### Hair-cell stimulation and functional imaging

Larvae at 5 and 6 dpf were prepared for GCaMP6s Ca^2+^ imaging as described previously ^58^. Primary posterior lateral-line neuromasts (NM1-NM4) were stimulated using a fluid jet. The fluid jet consisted of a pressure clamp (HSPC-1, ALA Scientific) attached to a glass pipette (inner tip diameter ∼ 50 μm). The glass pipette was filled with solution and used to deliver a 500-ms anterior and posterior mechanical stimulus. For GCaMP6s Ca^2+^ imaging a Bruker Swept-field confocal system with Prairie view software was used for image acquisition. The Bruker Swept-field confocal system was equipped with a Rolera EM-C2 CCD camera, and a Nikon CFI Fluor 60X/NA 1.0 water immersion objective. For calcium imaging, 5 plane Z-stacks were acquired every 0.5 μm (HC Apex calcium) or 1 μm (HC Base, pre- or post-synaptic calcium), at a 50 Hz frame rate, yielding a 10 Hz volume rate. Z-stacks were acquired using a piezoelectric motor (PICMA P-882.11-888.11 series, PI instruments) attached to the objective to allow rapid imaging along the Z-axis. 5 plane Z-stacks were projected into a single plane for further image processing and quantification. After post-synaptic GCaMP6s calcium measurements, larvae were labeled and with FM 4-64 (3 µM for 30 s) to label HCs.

After imaging, the raw images were registered to reduce movement artifacts. For pre- and post-synaptic GCaMP6s measurements, a circular region of interest (ROI) with a diameter ∼3 μm was placed on each synaptically active HC within a neuromast. For hair-bundle GCaMP6s measurements, a circular ROI with a ∼1.5 μm diameter was placed on the center of each individual bundle. After selecting an ROI, we calculated and plotted (ΔF/F_0_) within each ROI during the recording period. The signal magnitude was defined as the peak value of intensity change upon stimulation. Spatial heat maps to display calcium signals were generated as described previously ^58,59^. Heatmaps from anterior and posterior stimuli were combined into a single image for simplicity.

### Drug treatments and western blot analysis

Inhibition of lysosomal degradation was done using a previously published combination of pharmacological treatments ^41,42^. For this experiment, larvae were immobilized and mitochondria in axon terminals were photoconverted as described above. Immediately after photoconversion, larvae were removed from the agarose and placed in 10μm Pepstatin A (Fisher Scientific; BP26715)/10μm E64D (Enzo Life Sciences; BML-PI107-0001) in embryo media with 0.1% DMSO overnight before being imaged again the following day or lysed for protein extraction.

### Statistical analyses

For analyses of synapse number and size, statistical significance was determined by *t*-test with a Dunnett’s test to correct for multiple comparisons (synapse counts) or an unpaired *t*-test (areas).

For analyses of HC and afferent axon activity, statistical significance was determined by one-way ANOVA with a Dunnett’s test to correct for multiple comparisons.

For all other experiments, all statistical analyses were done using JMP14. For parametric analyses, ANOVAs were used with Tukey HSD post-hoc contrasts for pairwise comparisons. For non-parametric datasets, comparisons were done using Wilcoxon/Kruskal-Wallis analyses with corrected tests for each pair for multiple comparisons.

