## Supplemental Figures and resources for "Retrograde mitochondrial transport is essential for mitochondrial homeostasis in neurons"

Figure S1: Mitochondrial turnover from axon terminals is consistent across all conditions imaged. (**a**) Axon terminals in NM3 (mid-trunk neuromast) at 6 dpf show similar levels of mitochondria turnover to the same terminals 24 hrs after photoconversion at 4 dpf (see Fig. 1; n=23). (**b**,**c**) Terminal cluster neuromast axon terminals show similar levels of mitochondrial turnover after photoconversion at 4 (n=15) and 6 dpf (n=21). (**d**,**e**) Removing the larvae from their agarose chambers between imaging sessions shows similar loss of converted mitochondrial signal at both 4 (n=14) and 6 (n=15) dpf at NM3. (**f**) Motor neuron axons also show mitochondrial turnover within 24 hrs after photoconversion at 4 dpf (n=13). (**g**) Schematic of motor neuron axon photoconversion in the *Tg(neurod:mito-mEos)* transgenic line. (**h**) Images showing the conversion of mitochondria (magenta) and the loss of this signal 24 hrs after conversion in motor neuron axons. For all, ANOVAs with Tukey HSD posthoc contrasts.

Figure S2: Expression of PercevalHR in single motor neuron cell bodies and their axons at 4 dpf. (**a**,**b**) Single motor neuron cell body in *actr10*$^-$ mutant and wildtype sibling expressing PercevalHR. (**c**,**d**) Expression of PercevalHR in a single motor neuron axon in a wildtype (**c**) and *actr10*$^-$ mutants (**d**) larvae at 4 dpf. Asterisk on autofluorescent pigment cell. Scale bars – 10μm.

Figure S3: ATP SnFR analysis of cytosolic ATP levels at 4 dpf. (**a-d**) Expression of *5kbneurod:mRuby-ATPSnFR* in a single wildtype and *actr10*$^-$ mutant pLL neuronal cell body and associated axon terminal. (**e**) Schematic of the transient transgenesis used to express the mRuby tagged ATP SnFR in a single neuronal soma and its axon terminal in the pLL system. (**f**) Quantification of the ATP SnFR fluorescence intensity normalized to mRuby expression shows no difference in ATP levels between wildtype and *actr10*$^-$ mutant neurons (ANOVA). Sample size indicated on graph. Scale bars – 10μm.

Figure S4: Rescued *actr10*$^-$ mutant HCs effectively mechanotransduce upon stimulation but show inhibited synaptic responses. (**a**) Schematic of the neuromast indicating the Apex of the stereocilia where mechanotransduction takes place and the Base of the hair cell where the pre-synapse is located. (**b**,**c**) Wildtype and *actr10*$^-$ mutant HC rescue transgenic zebrafish larvae at 5 dpf show active mechanotransduction in HC stereocilia (Apex). (**b`**,**c`**) Heat map of Apex GCaMP6s fluorescence intensity changes with stimulation. (**d**,**e**) Quantification of the change in GCaMP6s fluorescence intensity showing active mechanotransduction in both wildtype and HC rescued *actr10*$^-$ mutants. (**f**,**g**) *actr10*$^-$ HC rescue mutants show reduced calcium influx at the presynapse. (**f`**,**g`**) Heat map of GCaMP6s fluorescence intensity changes at the HC base with stimulation. (**h**,**i**) Quantification of the change in GCaMP6s fluorescence intensity upon stimulation (*t*-test). Sample size indicated on graph. Scale bar – 10μm.

Figure S5: Immunolabeling of proteins identified in the mass spectrometry analysis as decreased in *actr10*$^-$ mutant mitochondria. (**a-d**) Immunofluorescence of ZZEF1 in the pLLg and axon terminals. (**e-h**) Immunofluorescence of Rps26 in the pLLg and axon terminals. (**i-p**) Immunolabeling of two proteins not altered in the mitochondrial mass spec data set, LamC and Aldh6a. Immunofluorescence signal in the pLLg was isolated from surrounding tissue using the ImageJ *Image calculator* tool for image preparation. Arrowheads point to a region of the axon terminal for each. Scale bar – 10μm.

### Figure S1

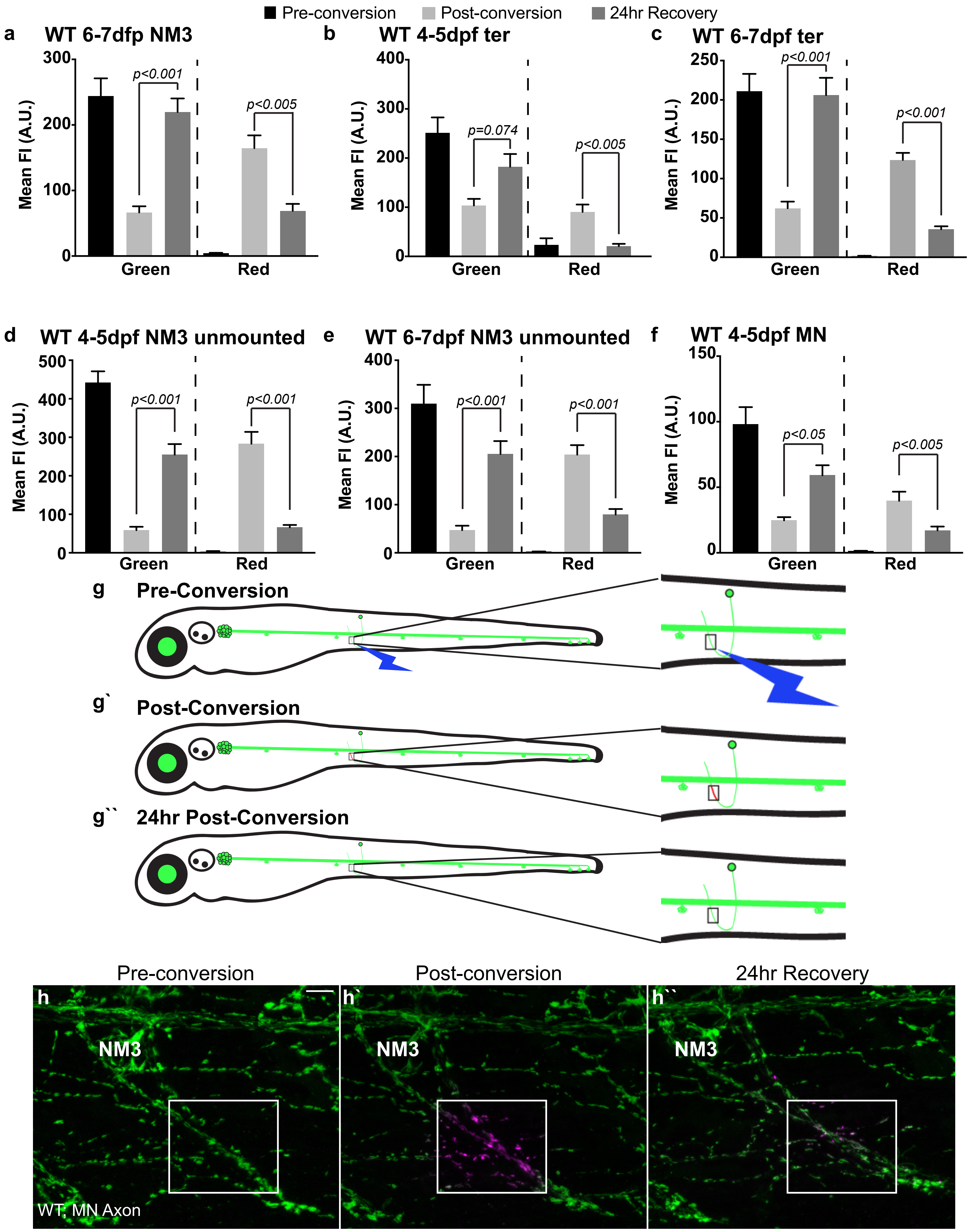

Pre-conversion
Post-conversion
24hr Recovery
a
WT 6-7dfp NM3
b
WT 4-5dpf ter
c
WT 6-7dpf ter
d
WT 4-5dpf NM3 unmounted
e
WT 6-7dpf NM3 unmounted
f
WT 4-5dpf MN
Mean FI (A.U.)
Green
Red
p<0.001
p<0.005
p=0.074
p<0.05
g
Pre-Conversion
g`
Post-Conversion
g``
24hr Post-Conversion
Pre-conversion
Post-conversion
24hr Recovery
h
h`
h``
NM3
WT; MN Axon

**Figure S2**

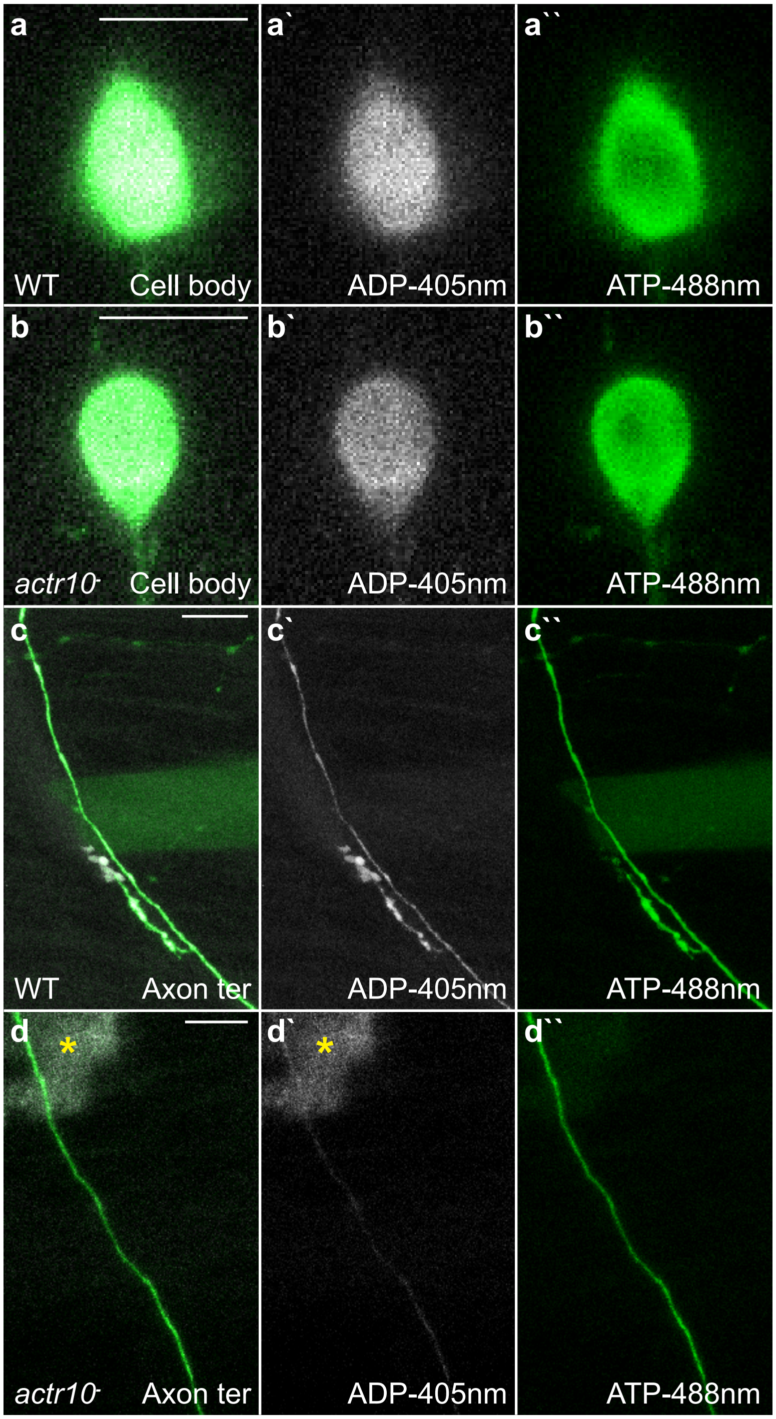
a
a`
a``
WT
Cell body
ADP-405nm
ATP-488nm
b
b`
b``
actr10-
Cell body
ADP-405nm
ATP-488nm
c
c`
c``
WT
Axon ter
ADP-405nm
ATP-488nm
d
d`
d``
actr10-
Axon ter
ADP-405nm
ATP-488nm

### Figure S3

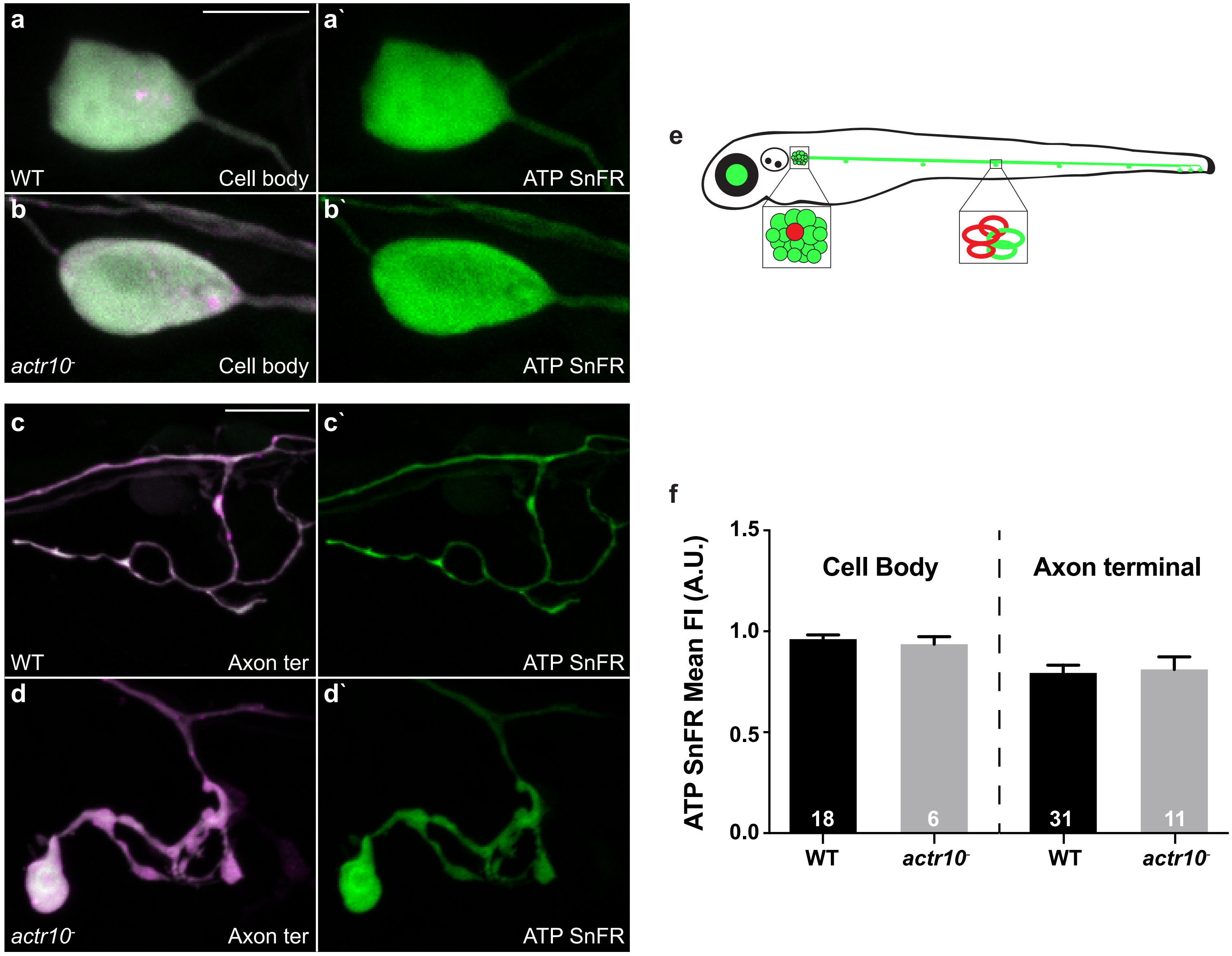

a
a`
WT
Cell body
ATP SnFR
b
b`
actr10-
Cell body
ATP SnFR
c
c`
WT
Axon ter
ATP SnFR
d
d`
actr10-
Axon ter
ATP SnFR
e
f
ATP SnFR Mean FI (A.U.)
1.5
1.0
0.5
0.0
Cell Body
Axon terminal
18
6
31
11
WT
actr10-
WT
actr10-

#### Figure S4

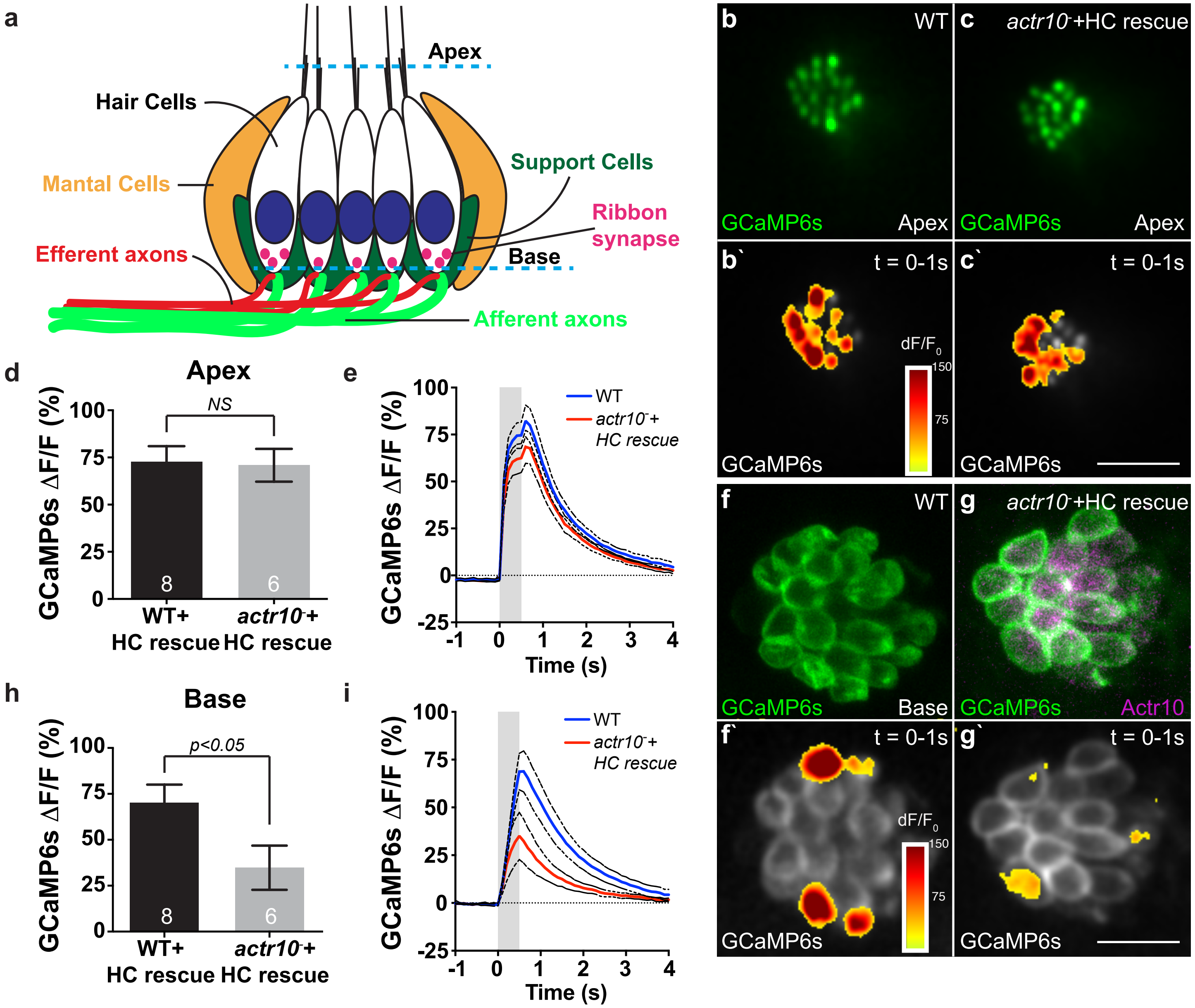

a
Apex
Hair Cells
Mantal Cells
Support Cells
Ribbon synapse
Efferent axons
Base
Afferent axons
b
WT
GCaMP6s
Apex
c
actr10⁻+HC rescue
GCaMP6s
Apex
b`
t = 0-1s
dF/F0
150
75
GCaMP6s
c`
t = 0-1s
GCaMP6s
d
Apex
GCaMP6s ΔF/F (%)
NS
WT+ HC rescue
actr10⁻+ HC rescue
8
6
e
GCaMP6s ΔF/F (%)
WT
actr10⁻+ HC rescue
Time (s)
f
WT
GCaMP6s
Base
g
actr10⁻+HC rescue
GCaMP6s
Actr10
f`
t = 0-1s
dF/F0
GCaMP6s
g`
t = 0-1s
GCaMP6s
h
Base
GCaMP6s ΔF/F (%)
p<0.05
WT+ HC rescue
actr10⁻+ HC rescue
i
GCaMP6s ΔF/F (%)
WT
actr10⁻+ HC rescue
Time (s)

#### Figure S5

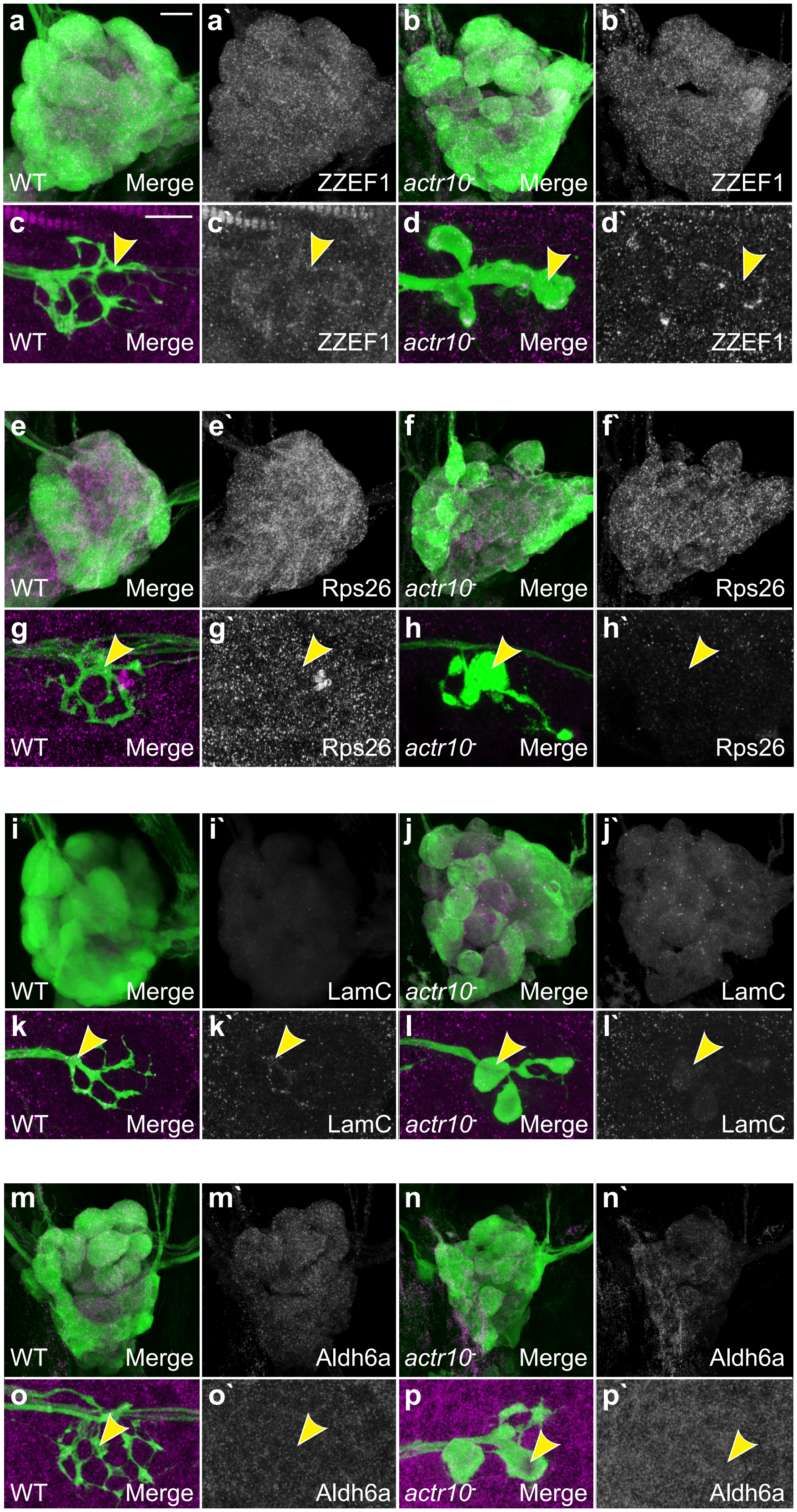

a
a`
b
b`
WT
Merge
ZZEF1
actr10-
Merge
ZZEF1
c
c`
d
d`
WT
Merge
ZZEF1
actr10-
Merge
ZZEF1
e
e`
f
f`
WT
Merge
Rps26
actr10-
Merge
Rps26
g
g`
h
h`
WT
Merge
Rps26
actr10-
Merge
Rps26
i
i`
j
j`
WT
Merge
LamC
actr10-
Merge
LamC
k
k`
l
l`
WT
Merge
LamC
actr10-
Merge
LamC
m
m`
n
n`
WT
Merge
Aldh6a
actr10-
Merge
Aldh6a
o
o`
p
p`
WT
Merge
Aldh6a
actr10-
Merge
Aldh6a

| Reagents or Resources | Source | Catalog number |
|---|---|---|
| Zebrafish Strains | | |
| AB | ZIRC | ZL1 |
| *actr10*$^{nl15}$ | (Drerup et al., 2017) | NA |
| *p150b*$^{nl16}$ | (Drerup et al., 2017) | NA |
| *TgBAC(neurod:egfp)*$^{nl1}$ | (Obholzer et al., 2008) | NA |
| *Tg(5kbneurod:mito-mEos)*$^{y568}$ | (Mandal et al., 2018) | NA |
| *Tg(5kbneurod:G-GECO)*$^{nl19}$ | (Mandal et al., 2018) | NA |
| *Tg(5kbneurod:mito-R-GECO)*$^{nl20}$ | (Mandal et al., 2018) | NA |
| *Tg(myo6b:mRFP-actr10)*$^{y610}$ | This paper | NA |
| *Tg(-6myo6b:GCaMP6s-CAAX)*$^{idc1Tg}$ | (Sheets et al., 2017) | NA |
| *Tg(hsp70l:GCaMP6s-CAAX-SiLL1)*$^{idc8Tg}$ | (Zhang et al., 2018) | NA |
| Antibodies for IF | | |
| GFP; 1:2000 | Aves | GFP-1020 |
| DsRed; 1:1000 | ThermoFisher | 632496 |
| Apex1; 1:100 | DSHB | CPTC-Apex1-2-s |
| ZZEF1; 1:500 | ThermoFisher | PA5-56856 |
| Sod2; 1:500 | ThermoFisher | PA1-31072 |
| RPS15a; 1:100 | ThermoFisher | PA5-24733 |
| RPS26; 1:500 | ThermoFisher | PA5-85689 |
| RPL27; 1:200 | ThermoFisher | PA5-51728 |
| Tom20; 1:200 | Santa Cruz | sc-11415 |
| Ribeye b; 1:10,000 | (Sheets et al., 2011) | NA |
| MAGUK; 1:500 | NeuroMab | K28/86 |
| Myosin7a; 1:1000 | Proteus Biosciences | 25-6790 |
| AlexFluor 488/568/647; 1:1000 | ThermoFisher | assorted |
| Antibodies for Western blot | | |
| LC3; 1:2000 | Novus | NB100-2331SS |
| GFP; 1:2500 | ThermoFisher | A11122 |
| Anti-rabbit*HRP | Jackson Immunoresearch | 711-036-152 |
| Drug Treatments | | |
| Pepstatin A; 10$\mu$m | Fisher Scientific | BP26715 |
| E-64-D; 10$\mu$m | Enzo Life Sciences | BML-PI107-0001 |
| Vital Dyes | | |
| Tetramethylrhodamine ethyl ester (TMRE) | ThermoFisher | T669 |
| FM1-43 | ThermoFisher | T35356 |

| Genotyping target | Forward Primer | Reverse Primer |
|---|---|---|
| *actr10*$^{nl15}$ | TGTTTTCGGATGAACTGCCTG | CGTGTAGGCCGCTCCTAAAT |
| *p150a* | TAGTGTGCAGATCCAATATGGC | ATTTCCCAGAGGCGAAGAGT |
| *p150b*$^{nl16}$ | TCGGCTTCTGCAGGAGAGAT | CTGGGTCCCGGGATTGGAG |
| *Tg(myo6b:mRFP-actr10)*$^{y610}$ | CGCCTACAAGACCGACATCA | TGGACATCAGGTGACTGGGA |

| Construct | Citation | Sensor original source |
|---|---|---|
| *5kbneurod:mito-TagRFP* | (Drerup et al., 2017) | (Fang et al., 2012) |
| *5kbneurod:mito-TIMER* | (Mandal et al., 2018) | Addgene: 50547 |
| *5kbneurod:PercevalHR-mCherryCAAX* | (Mandal et al., 2018) | Addgene: 49082 |

| | | |
|---|---|---|
| *5kbneurod:roGFP2* | This paper | Addgene: 64977 |
| *5kbneurod:mito-roGFP2* | This paper | Addgene: 64977 |
| *5kbneurod:mRuby-ATPSnFR* | This paper | Lobas et al 2019 |
| *mnx1:PercevalHR-mCherryCAAX* | This paper | Addgene: 49082 |

| **shRNA against Rat Actr10** | **Sequence** |
|---|---|
| #1 | ggcatgtctaagccgatcaaa |
| #2 | ggtcctggattgtggatatag |
| #3 | gtgcttggatcaatcagagat |
